# DNA spike-ins enable confident interpretation of SARS-CoV-2 genomic data from amplicon-based sequencing

**DOI:** 10.1101/2021.03.16.435654

**Authors:** Kim A. Lagerborg, Erica Normandin, Matthew R. Bauer, Gordon Adams, Katherine Figueroa, Christine Loreth, Adrianne Gladden-Young, Bennett Shaw, Leah Pearlman, Erica S. Shenoy, David Hooper, Virginia M. Pierce, Kimon C. Zachary, Daniel J. Park, Bronwyn L. MacInnis, Jacob E. Lemieux, Pardis C. Sabeti, Steven K Reilly, Katherine J. Siddle

**Affiliations:** Broad Institute of Harvard and MIT, 415 Main Street, Cambridge, MA 02142, USA; Harvard Program in Biological and Biomedical Sciences, Harvard Medical School, Boston, MA 02115, USA; Department of Systems Biology, Harvard Medical School, Boston, MA, USA; Division of Infectious Diseases, Massachusetts General Hospital, Boston, MA, USA; Department of Pathology, Massachusetts General Hospital, Boston, MA, USA; Pediatric Infectious Disease Unit, MassGeneral Hospital for Children, Boston, MA, USA; Department of Pathology, Harvard Medical School, Boston, MA, USA; Department of Medicine, Harvard Medical School, Boston, MA 02115, USA; Infection Control Unit, Massachusetts General Hospital, Boston, MA, USA; Department of Immunology and Infectious Diseases, Harvard T. H. Chan School of Public Health, Harvard University, Boston, MA, USA; Massachusetts Consortium on Pathogen Readiness, Boston, MA 02115, USA; Howard Hughes Medical Institute, 4000 Jones Bridge Rd, Chevy Chase, MD 20815, USA

## Abstract

The rapid global spread and continued evolution of SARS-CoV-2 has highlighted an unprecedented need for viral genomic surveillance and clinical viral sequencing. Amplicon-based sequencing methods provide a sensitive, low-cost and rapid approach but suffer a high potential for contamination, which can undermine lab processes and results. This challenge will only increase with expanding global production of sequences by diverse research groups for epidemiological and clinical interpretation. We present an approach which uses synthetic DNA spike-ins (SDSIs) to track samples and detect inter-sample contamination through a sequencing workflow. Applying this approach to the ARTIC Consortium’s amplicon design, we define a series of best practices for Illumina-based sequencing and provide a detailed characterization of approaches to increase sensitivity for low-viral load samples incorporating the SDSIs. We demonstrate the utility and efficiency of the SDSI method amidst a real-time investigation of a suspected hospital cluster of SARS-CoV-2 cases.

## Introduction

The COVID-19 pandemic has demonstrated the need for sensitive, fast, and low-cost viral genomic sequencing in labs distributed throughout the world. Genome sequencing early in the outbreak allowed for rapid identification of SARS-CoV-2 and enabled implementation of diagnostics in many countries. In the year since, an ongoing scaling up of genomic data has provided new insights into the diversity, evolution and transmission of the virus, which has increasingly been used to guide important public health interventions. In particular, viral genome sequencing has been used to characterize the epidemiology of clusters and superspreading events ^1–3^. Concurrently, genome sequencing to monitor the emergence of new lineages and the spread of variants of concern (VoC) has been a of global priority ^4^. As clinically relevant characteristics of COVID-19 for each VoC become more well understood, such as therapeutic evasion, the role of genomic sequencing in the clinical setting will only grow.

Multiplexed amplicon-based genome sequencing methods have accelerated the unprecedented scale of SARS-CoV-2 genomic surveillance due to improved sensitivity, speed and cost over unbiased, low-amplification RNA sequencing approaches ^5^. In just a year since the first genome sequence enabled the identification of SARS-CoV-2, hundreds of thousands of complete genomes have been released by several hundred laboratories, the vast majority (over 90% of Short Read Archive submissions), using amplicon-based approaches that target the virus’ genome for amplification and subsequent sequencing. An open-access tiled primer set developed by the ARTIC network (https://artic.network/) is the most widely used method for SARS-CoV-2 specific genome amplification followed by sequencing on either Illumina or nanopore instruments ^6,7^. A wide array of protocols and publications that integrate these ARTIC primers with different amplification and library construction indexing strategies are now available ^8,9^.

However, by the very nature of using approaches that rely on high amplification of viral genomes, contamination is a critical risk faced by laboratories processing these samples. The 35 or more cycles of virus-specific PCR produce trillions of SARS-CoV-2 amplicons in a single reaction, some of which may end up in the laboratory environment via aerosolization, potentially confounding studies where viral detection is sensitive to only tens of molecules ^10,11^. Many labs performing viral sequencing are often processing multiple large batches (96) of samples in parallel, further increasing chances of direct cross contamination or sample swapping ^12^. Moreover, as SARS-CoV-2 has relatively low genetic diversity and high superspreading potential ^13,14^, many genomes are expected to be identical at the consensus level, a pattern that could also be observed due to contamination ^11,15–17^. Such sample swaps or contamination could confound phylogenetic identification of clusters of infections that may represent transmission events or lead to less effective treatment for a mis-identified clinically relevant variant; thus, mitigating this risk is critical.

To meet the genomic surveillance goals laid out by local and world governments, sequencing efforts are being scaled to thousands of centers, many performing viral genomics for the first time ^18,19^. Additional laboratories will enter the SARS-CoV-2 sequencing space with an emphasis to rapidly surveil VoCs for clinical significance, necessitating stringent requirements to ensure the integrity of SARS-CoV-2 genomes being produced. This rapid expansion of genomic data generation for surveillance, the emerging value of sequencing for clinical decision making, and the high potential for contamination in amplicon-sequencing, makes clear the urgent need for new tools to track samples with utmost precision and accuracy. While inclusion of internal standards are commonplace for mass spectrometry based assays in proteomics and metabolomics ^20,21^, and nucleic acid spike-ins are often used to detect sample swaps and contamination for RNA sequencing ^22^, there are no readily available and widely used quality control methods for amplicon-based genomic sequencing. While some technical assay controls exist ^23^, to date, there are no established control measures to counter for contamination risk in SARS-CoV-2 sequencing.

Here we designed, optimized, and implemented a novel sample identification method using synthetic DNA spike-ins (SDSIs) that is broadly compatible with SARS-CoV-2 sequencing approaches and settings. We implemented these SDSIs for Illumina sequencing with SARS-CoV-2 specific amplification using the ARTIC consortium’s primer panel designs. To maximize recovery of consensus genomes from samples with low viral loads, we compared improvements to key amplification and library construction steps and validate the accuracy of the assembled genomes. We propose a modified protocol, hereafter termed SDSI+ARTIC, that provides increased confidence in the veracity of genomes with minimal extra cost and time that can be applied to epidemiological and clinical investigations of SARS-CoV-2 (**Fig 1**).

**Figure 1.**
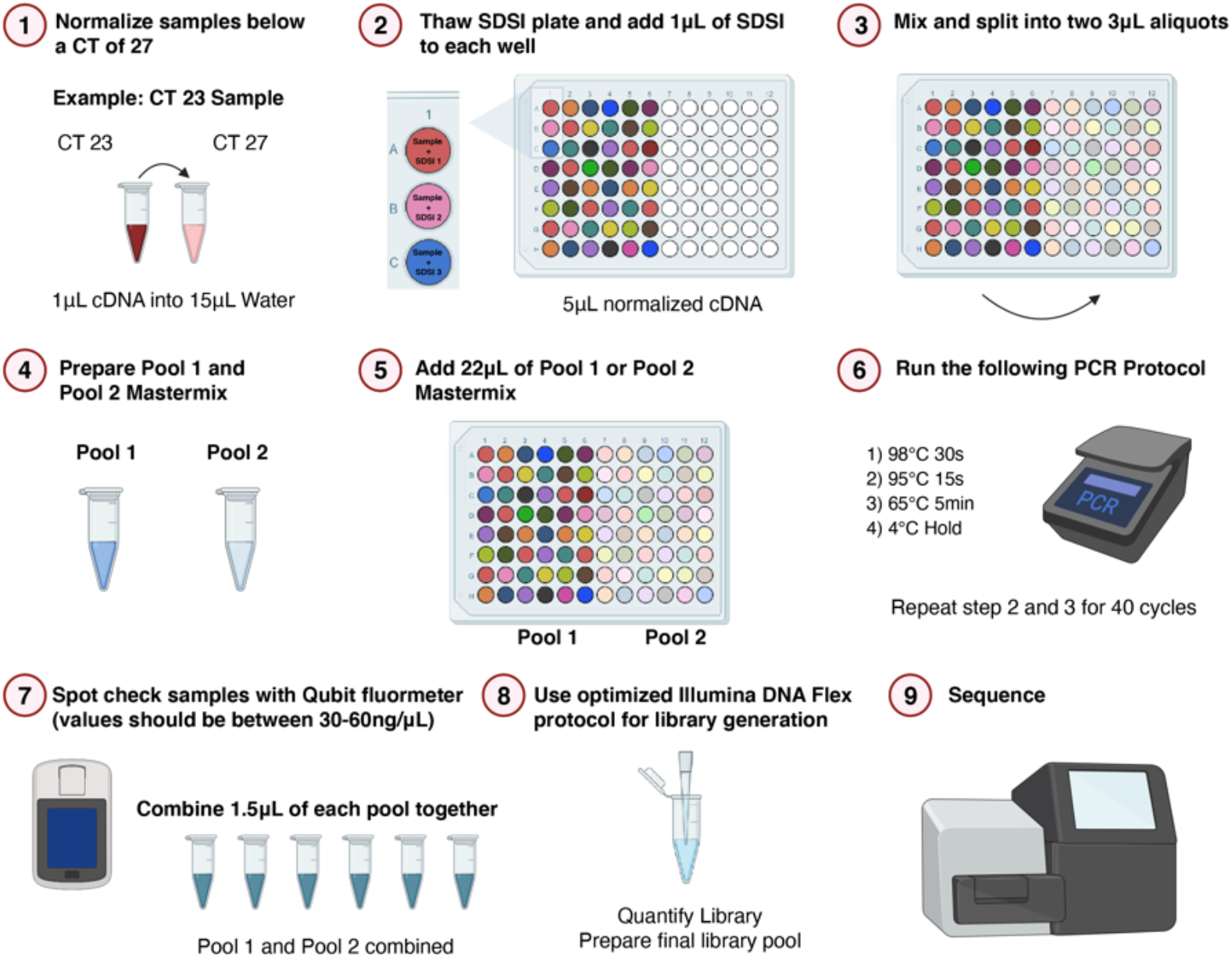
SDSI+ARTIC Amplicon-Sequencing Protocol. Illustrative workflow for 48 samples through the SDSI+ARTIC amplicon-sequencing pipeline. A unique, synthetic DNA spike-ins (SDSI) will be added to each sample to allow for contamination tracking and accurate sample identification in analysis. Created with BioRender.com.

## Results

### Design and in silico validation of synthetic DNA spike-ins for amplicon-based sequencing

In designing a system for sample tracking and contamination tracing that would be applicable to a variety of viral amplicon-based sequencing strategies, we investigated the use of SDSIs that consisted of a core uniquely identifiable sequence flanked by constant priming regions. With such a design, a single additional primer set could be integrated into a multiplexed PCR to co-amplify any SDSI with the primary reaction target(s) **(Fig 2a)**. To enable maximum flexibility of use, these primers must be compatible with a wide variety of PCR reactions and highly specific for amplifying SDSIs, and the SDSI amplicons ought to amplify at similar rates to each other and primary reaction target(s). To be most sensitive in detecting potential contamination, and avoid false positive results, the core, unique SDSI sequences should be sufficiently distinct from one another, as well as sequences commonly found in laboratories. Such a design would enable in-sample labeling, where different amplified samples processed together could be associated reliably with unique SDSIs. Not only would this provide a sample-specific internal control that enables identification of sample swaps, but the association between SDSI and amplified sample content would also illuminate viral sequence contamination with high resolution and accuracy.

**Figure 2.**
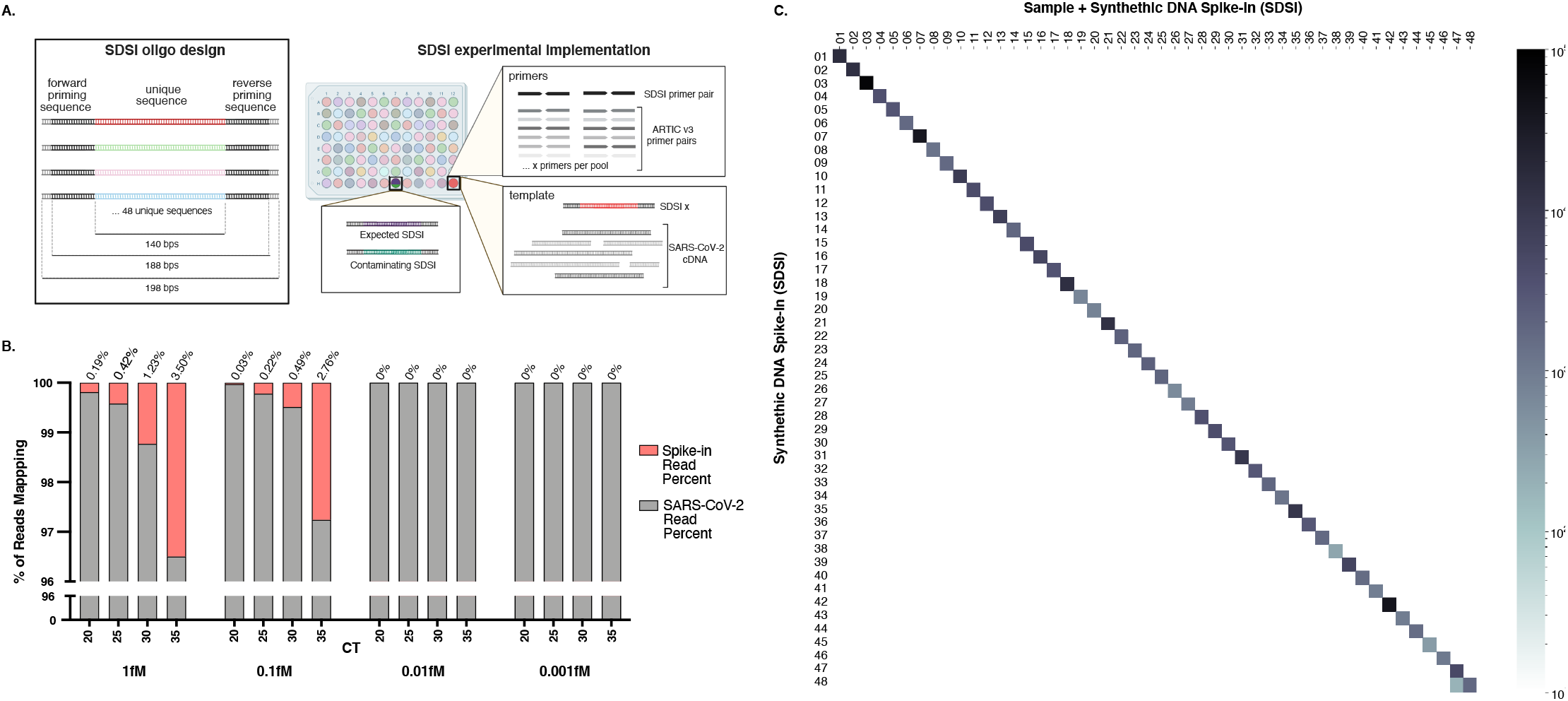
Synthetic DNA oligos spiked into amp-seq reactions flag contamination and sample swaps. **A.** Schematic of SDSI design. Each oligo contains 140 bp of unique sequence flanked by common primer binding sites. Primers designed to amplify all SDSIs are added to ARTIC primer pools, and a unique SDSI is added to each clinical sample. Identification of multiple SDSIs in the same sample indicates contamination. Created with BioRender.com. **B.** In a titration of SDSIs across clinical samples with variable CTs, the number of reads mapping to both SARS-CoV-2 and the SDSI were quantified, and the percentage of each was calculated. **C.** For each of 48 unique clinical samples (on the horizontal axis), reads mapping to each of 48 unique SDSIs (on the vertical axis) were quantified (log read count). Samples and SDSIs were ordered such that the intended match is on the diagonal of this matrix, thus any off-diagonal signal would reveal non-specific identification of SDSIs or contamination of SDSIs across samples.

To meet the requirement of reliable SDSI identification, we selected 48 distinct DNA sequences from the genomes of diverse, uncommon archaea to serve as the core portion of the SDSIs, precluding false detection and cross-identification. Since false detection of an SDSI would occur if its sequence shared significant homology with other genetic material in a sample, we based these sequences on archaea, which are divergent from organisms found in typical laboratory or clinical settings **(Sup Table 1)**. A permissive search performed against the entire NCBI database confirmed that 43/48 SDSI sequences had significant homology (>75% sequence identity over >75% query cover) exclusively within the domain archaea; the remaining 5 SDSIs had homology to a handful of rare bacterial genera unlikely to be found in laboratories (**Sup Table 1**). While this limited homology outside of the domain archaea maximized the potential for broad applications, we also specifically verified that each core SDSI sequence was unlikely to be confused with the expected COVID-19 clinical sample content and confirmed that all sequences had no homology (>50% sequence identity over >50% query cover) with either *Homo sapiens* or SARS-CoV-2. Each SDSIs consisted of a 140 bp stretch of variable sequence. We confirmed that all SDSIs were significantly different from each other to prevent misidentification; among all SDSIs, the minimum pairwise Hamming distances of the 140 bp stretch of unique sequence was 84 (mean=105; max=121).

We further considered design specifications to enable specific, consistent amplification of SDSIs in multiplexed PCR reactions. We designed an SDSI primer pair (and corresponding priming regions) that had limited homology to common organisms in order to preclude off-target priming in the PCR reaction that could outcompete amplification of a primary target. The primer pair also had a common length (24 bps) and GC content (45.8%) further promoting their compatibility with many multiplexed PCR reactions, including SARS-CoV-2 amplicon sequencing strategies (https://artic.network/). Since each SDSI was identically sized and shared a priming region, a similar amplification rate was expected across all SDSIs. Across the entire SDSI amplicons, we avoided extremes of GC content (range: 35-65%) in order to promote similar amplification rates across different SDSIs and to viral amplicons (e.g., the GC content of the SARS-CoV-2 genome is roughly 37±5%) ^24^ (**Sup Fig 1a**).

### Application of SDSIs to SARS-CoV-2 sequencing

The addition of SDSIs into the ARTIC multiplexed PCR provided a sample-specific internal control and did not detrimentally affect the amplification of SARS-CoV-2 cDNA. The SDSI primers did not produce any nonspecific amplification, including in the presence of cDNA from a nasopharyngeal swab sample, supporting the expectation that primers shared limited homology with genomic material from clinical samples **(Sup Fig 1b)**. All SDSIs amplified in an ARTIC SARS-CoV-2 PCR reaction with SDSI primers included, in each case yielding a single clean product of the expected size **(Sup Fig 1c)**. To prevent SDSIs from overtaking the amplification and sequencing of SARS-CoV-2 amplicons, we optimized the amount of SDSI added to each reaction through limited titration (**Fig 2b; Sup Fig 2**). Through a dilution series we found that 1μl of a 1fM SDSI resulted in the reliable detection of the SDSI across a range of CT values (CT 20, 25, 30, 35) while the majority of reads (>96%) still mapped to SARS-CoV-2 with no apparent alteration in coverage across the genome **(Fig 2b; Sup Table 2)**.

We performed SDSI+ARTIC sequencing on a set of 48 SARS-CoV-2+ clinical samples to demonstrate its feasibility for tracking samples in a large batch. After adding a different SDSI to each sample, we found that 47/48 SDSIs were identified exclusively in the anticipated sample, validating the use of SDSIs as an internal control and for identifying sample swaps **(Fig 2c)**. While not yet formally tested, we expect that in the case of within-batch contamination, SDSIs associated with contaminating samples would be identified in samples where they were not expected, as was the case with SDSI_48 which we detected in the intended sample, as well as a neighboring sample in the batch **(Fig 2a, 2c)**. In Sample 47, 96% of reads mapping to all SDSIs mapped to the expected SDSI (SDSI_47) while 4% mapped to SDSI_48. We recommend manual curation of genomes assembled from any sample with <99% of SDSI reads mapping to the expected SDSI, and therefore compared the genome from Sample 47 to that of the potentially contaminating sample (Sample 48). Ultimately no further genome validation was warranted because we did not assemble a genome from Sample 47 and it was removed from further analyses. Interestingly, assembled portions of the genome revealed two single nucleotide variations (SNVs) that differentiated it from the genome from Sample 48, which indicates that viral reads did not solely originate from contamination. Nevertheless, this case reveals the potential prevalence of undetected contamination and underscores the importance of a method for identifying it.

### Improving genome recovery and coverage for SARS-CoV-2 Viral Genomics

There are a number of other critical technical enhancements that can further increase the quality of genomic data we can generate for epidemiological and clinical investigations. Clinical samples can have a wide range of viral loads and/or varying degrees of sample degradation, necessitating improved methods for capturing complete genomes from samples with lower RNA amounts and/or degraded viral RNA. As these samples often poorly amplify, they can also result in uneven genome coverage. Additionally, scaling sequencing efforts across a wide spectrum of samples requires a flexible method to capture low quality samples while balancing the excessive amplification of from higher viral titers, in which the latter increases the risk of contamination, confounding accurate interpretation of sequencing data.

To better capture genomes from samples with lower RNA amounts or degraded viral RNA we tested various modifications to cDNA generation. To increase the recovery of such genomes, we examined a number of reverse transcriptases and found more processive ones yielded more and longer cDNA generation. Comparing cDNA produced with Superscripts III, IV, or IV-VILO across a range of clinical viral loads (high viral load: CT <20, mid-high viral load: CT 20-25, mid-low viral load: CT 25-30, and low viral load: CT >30), SSIV-VILO and SSIV produced the highest number of amplicons, with at least 10X coverage across 13 samples (SSIII: 72.64%, SSIV: 81.93%, SSIV-VILO: 86.97%) **(Fig 3a)**. These processive reverse transcriptases also displayed lower variability as measured by the percent of amplicons with <20% mean coverage (SSIII: 36.89% SSIV: 31.24% SSIV-VILO: 22.45%) **(Sup Fig 4a)**. In our hands, generation of longer and higher quality cDNA was an essential prerequisite for successful amplification of the SARS-CoV-2 genomes needed for subsequent sequencing.

**Figure 3.**
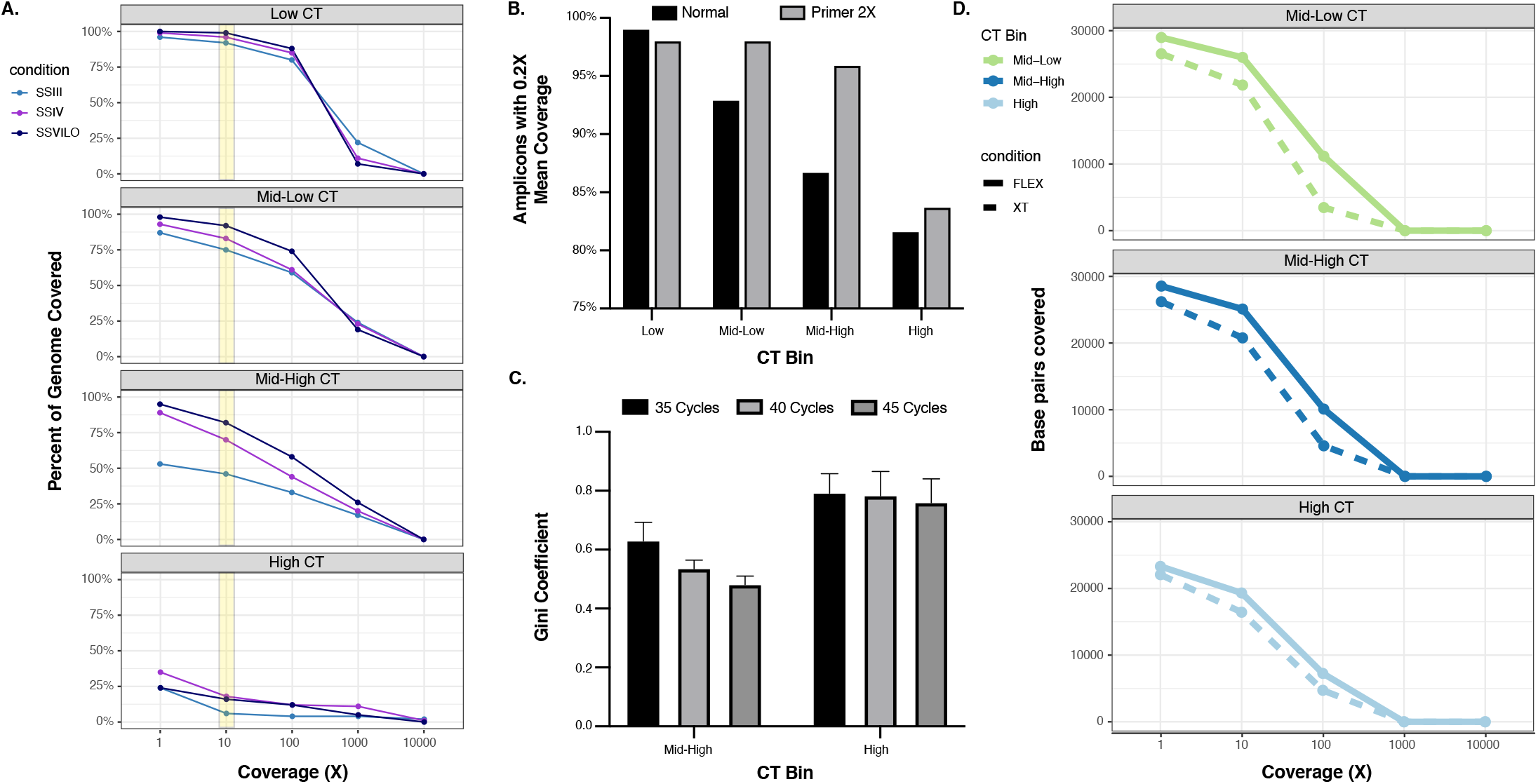
Maximizing Genome Recovery and Coverage with SDSI+ARTIC. **A.** The percent of the target genome covered at various depths of coverage with various reverse transcriptases, used for cDNA synthesis. Data represents four individual samples. Yellow bar highlights comparison between the reverse transcriptases a coverage depth of 10X. **B.** Amplicons with at least 0.2X of the mean amplicon coverage with the normal ARTIC v3 primer pools or with a modified primer pool with a 2X concentration of 20 poor-performing ARTIC primer pairs. Four samples with low, mid-low, mid-high, and high CTs were used. **C.** Gini coefficients for two mid-high CT samples and four high CT samples when using either 35, 40, or 45 cycles for the ARTIC PCR. Error bars represent standard deviation. **D.** Comparison of Nextera DNA Flex and Nextera XT on the number of SARS-CoV-2 base pairs covered at various depths of coverage for three samples at different CTs.

We then tested a number of modifications to the ARTIC PCR reaction that could increase overall amplification and multiplexed amplicon uniformity, starting with different primer concentrations and DNA polymerases, to aid in the recovery of complete genomes with the smallest number of reads. We found that increasing (2x) primer concentrations (20.8nM final) for low efficiency amplicons increased coverage in these amplicons, even enabling whole genome recovery for multiple samples **(Fig 3b; Sup Fig 5; Sup Table 3).** By testing five DNA polymerases and conditions under standard ARTIC PCR conditions (**Methods**), we found Q5 Hot Start High-Fidelity 2x Master Mix and KAPA reactions yielded the highest amplification (average 85.3nM and 56nM respectively) (**Sup Fig 4b**). In turn, this suggested methods that produce the highest amplification across a range of viral titers yield more uniform coverage and reduce read depth required for downstream sequencing.

We then explored the effects of different numbers of PCR cycles, DNA-hybridization steps, temperature ramp speeds, and primer design. Attempting to recover low viral load samples by increasing the number of PCR cycles, we found greater coverage uniformity with increasing cycles (**Fig 3c**). However, at 45 cycles 3 SNVs that were not present in lower-amplified samples were noted. In balancing increased amplification from low viral load samples with the potential increase in erroneous SNV calls with increased cycling we implemented a 40 cycle PCR. Additional modifications such as DNA-rehybridization steps ^25^ or slower temperature ramp speeds had no significant effects **(Sup Fig 4c, 4d)**. We found that an alternative primer design using shorter (~150bp) amplicons, the Paragon Genomics CleanPlex SARS-CoV-2 panel, did recover more complete genomes in very low viral load samples (CT >35), but had lower accuracy and drops in coverage even in high viral load samples which resulted in missed SNV calls (**Sup Fig 3a, 3b**), consistent with previous reports ^12,26^. In our hands, a 40 cycle PCR utilizing ARTIC primers without alterations to the ramp temperatures or speeds provided the greatest amplification without deleterious effects of increasing unfounded SNV calls.

In achieving more uniform genome coverage in samples with lower viral loads with increased cycling, higher viral load samples will amplify much greater than the amount required for sequencing. To mitigate the risk of contamination from such highly amplified libraries, we found that scaling down (.5X) Illumina DNA Flex library construction reagents provide a limit on amplified material. Notably, this limitation did not impact final library size distributions, while having the desired effect of generating final sequencing libraries at half their original concentrations. This approach also had the added benefit of nearly halving the library construction cost per sample (**Sup Table 4; Sup Table 5**). In our comparisons, we also observed the Nextera DNA Flex generated greater coverage depth and uniformity than DNA XT, further supporting its use in Illumina-based ARTIC sequencing (**Sup Fig 6**). The degree of overamplification of high viral load samples compelled us to reduce the overall DNA Flex reactions in efforts to balance appropriate library yield for sequencing and contamination.

### SDSI+ARTIC sequencing benchmarks well against unbiased sequencing

We observed near perfect sequence concordance when comparing SDSI+ARTIC to unbiased sequencing, which has served as the gold standard for generating error-free viral genomes and for capturing divergent SARS-CoV-2 strains ^5,12^. Our comparisons of these two sequencing approaches is consistent with reports from other groups that have demonstrated that ARTIC sequencing improves sensitivity while maintaining a high level of concordance at the consensus genome level ^12^. As the inclusion of SDSIs to the standard protocol slightly reduced the sequencing depth for SARS-CoV-2 and could affect the efficiency of the ARTIC PCR and/or library construction, we further compared the modified SDSI+ARTIC method to unbiased sequencing (**Fig 2b**). We sought to perform this comparison on a large batch of samples since previous analyses of targeted enrichment approaches and unbiased sequencing were limited by small sample sets (<24 samples) which may not recapitulate the full extent of contamination risk and performance across a wider range of samples ^12,27^.

To serve as a direct comparison, we performed SDSI+ARTIC method on a batch of 96 samples (89 unique patient samples + 7 water controls) that were previously sequenced using an unbiased metagenomic sequencing approach ^1^. The 89 patient samples consisted of diverse viral lineages and a broad range of viral loads (CT range = 11.9-37.4; mean = 27.4) (**Sup Fig 7a**). As expected, SDSI+ARTIC outperformed unbiased sequencing in the number of complete genomes (>98% assembled) and partial genomes (>80% assembled) (**Fig 4a,4b**; **Sup Fig 7b).** We assessed coverage uniformity in both methods, as increasing uniformity reduces the sequencing depth required to generate reliable genomes, thus improving throughput and efficiency ^28^. Comparisons of Gini coefficient for each sample that generated an assembly revealed that unbiased sequencing had more uniform coverage up to a CT of 25 (N=31, Gini Coeff = 0.240 ± 0.046 (unbiased) vs 0.428 ± 0.026 (SDSI+ARTIC)), while SDSI+ARTIC generated more uniform coverage for samples above a CT of 25 and below CT 37 (N=39, Gini Coeff = 0.766 ± 0.265 (unbiased) vs 0.554 ± 0.124 (SDSI+ARTIC)) (**Sup Fig 7c)**. As expected, both methods yielded less uniform coverage in samples with lower viral loads.

**Figure 4.**
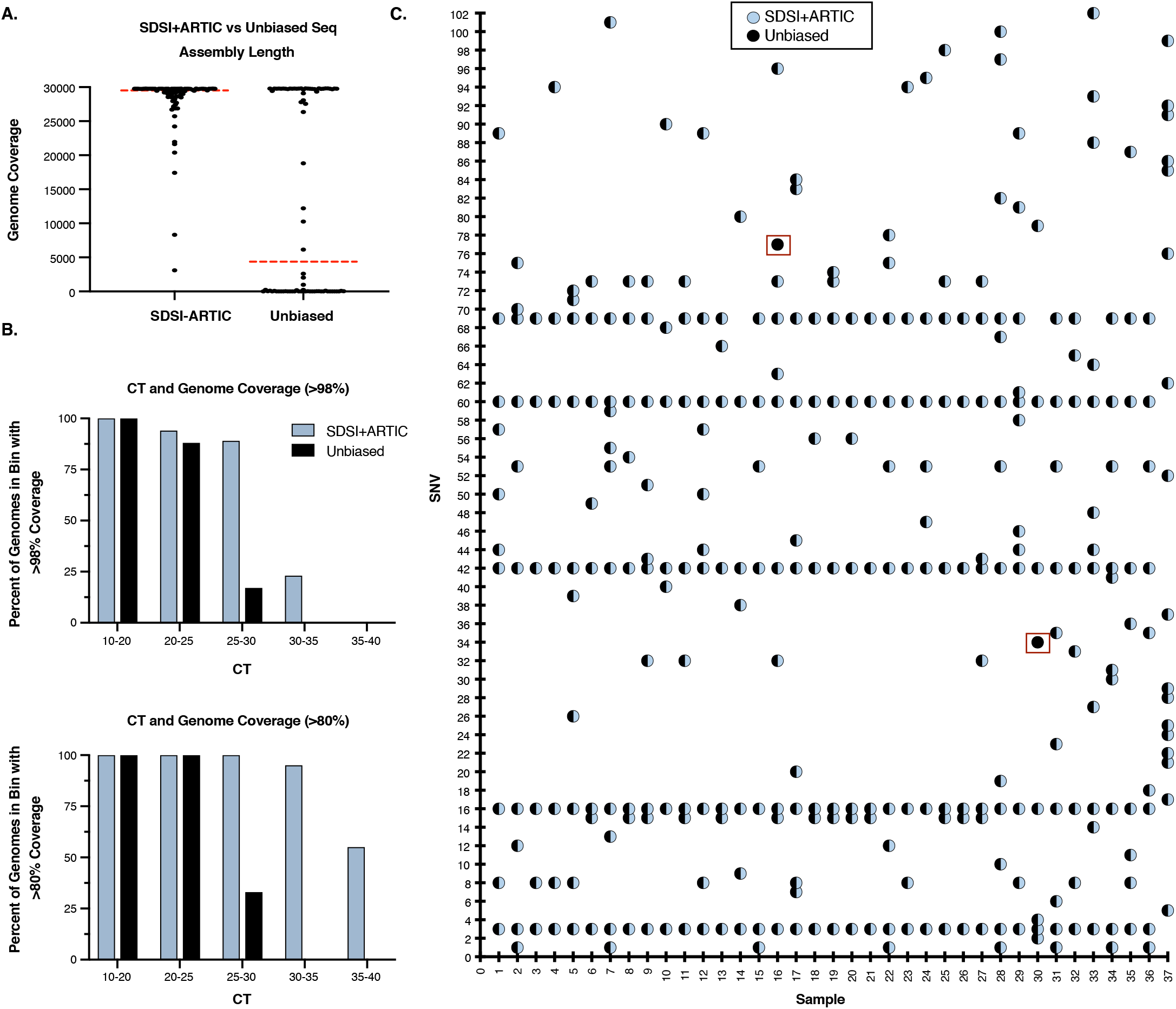
SDSI+ARTIC has a high level of concordance with unbiased sequencing across a wide range of clinical CTs. **A.** SDSI+ARTIC (N=81) and unbiased sequencing (N=81) assembly lengths. All samples were downsampled to 975,000 reads. Dotted red line indicates median assembly length (SDSI+ARTIC = 29,577; Unbiased = 4,389). **B.** Percent of assemblies with greater than 98% or 80% coverage in different CT bins (SDSI+ARTIC N=81; Unbiased N=81) (downsampled to 975,000 reads). **C.** SNV concordance plot between SDSI+ARTIC and unbiased consensus sequences. Two discordant SNVs, outlined in a red box, were found.

Most importantly, SDSI+ARTIC displayed high concordance in sequence variant identification, producing only two divergent SNV calls out of 332 total SNVs identified compared to the reference (Wuhan-Hu-1) **(Fig 4c**). To make the most equivalent comparison, we compared consensus sequences without downsampling using only samples that produced a full genome in both methods (N=37) (**Methods**). The discordant SNVs, which matched the reference sequence, were observed in two samples, and occurred in different regions of the viral genome. Both positions fell in ARTIC primer regions and matched the primer sequence even though primer trimming was performed and confirmed by manual inspection. Additionally, the coverage depth in the regions of the SNVs was greater than 1000X for both platforms in both samples and the SNVs persisted with both relaxed (n=3) and conservative (n=20) minimum coverage thresholds. We believe one of these discrepancies likely arose during the ARTIC PCR, whereas manual inspection of the other position (a C9565T mutation in unbiased sequencing) indicated the presence of intra-host variation in both methods with a variant allele frequency of 39.4% (SDSI+ARTIC) and 59.2% (unbiased sequencing). Overall, the discordance rate between SNV calling for SDSI+ARTIC and unbiased sequencing was 0.6%. When comparing concordance between the methods across all nucleotides (29,728 bp covered by the ARTIC panel after primer trimming), the concordance rate among 35 of the samples was 100% with the two aforementioned samples having a concordance rate of 99.997%. Notably, these two mismatches did not result in lineage misassignment for either sample.

### SDSI+ARTIC sequencing confirms a suspected nosocomial cluster

To underscore the real-world value of using SDSI+ARTIC to generate high-confidence genomes, we applied our method to investigate a putative SARS-CoV-2 cluster from Massachusetts General Hospital (MGH) for which the Infection Control Unit suspected nosocomial transmission. Viral sequence variation can distinguish between a common source of infection, characterized by many identical or highly genetically similar sequences, and independently acquired infections, that are expected to be genetically distinct. Identification of multiple identical genomes within a batch can also call into question the validity of such findings, which is challenging to rule out given standard amplicon-based sequencing workflows. Therefore, cluster investigations which aim to confirm or refute transmission events based on viral sequence would greatly benefit from SDSIs to ensure they exclude laboratory contamination. To this end, we sequenced 22 samples with SDSI+ARTIC; 14 samples suspected to be part of the cluster based on epidemiological contact-tracing, and 8 unlinked samples as controls.

The SDSI+ARTIC method enabled efficient and confident identification of a cluster of infections, with samples processed within 24 hours and final genomes assembled within 52 hours of biosample receipt. We assembled 17 genomes (>80% complete), and samples that did not yield a complete genome were those with lower viral loads (CT > 30). Of the 11 samples that we assembled genomes from that were suspected to be part of the cluster, 10 were genetically highly similar (0-1 consensus nucleotide difference) (**Fig 5a)** and a phylogenetic tree of these sequences alongside >4000 other genome sequences showed that these samples were distinct from other samples from Massachusetts around the same time **(Fig 5b)**, strongly suggesting that this cluster did arise from nosocomial transmission. One sample, MA-MGH-02834, differed from other putative cluster-associated samples by 18-19 consensus-level variants, suggesting that this infection was likely acquired independently of the nosocomial cluster.

**Figure 5.**
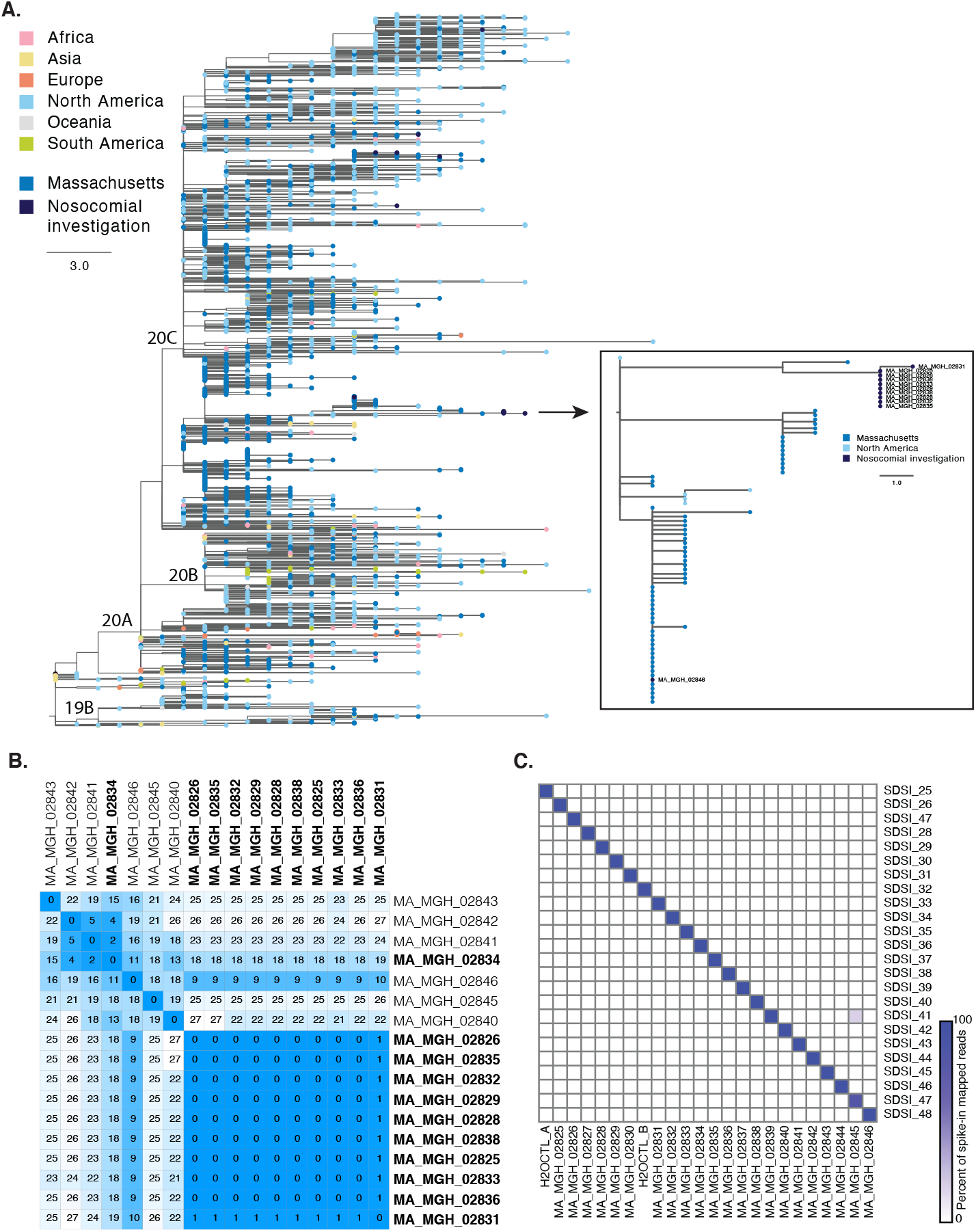
Deployment of SDSI+ARTIC to assess for possible nosocomial transmission. **A.** Phylogenetic tree showing the location of the putative cluster sequences in the context of a global subset of circulating SARS-CoV-2 diversity. Zoom box shows the 10 highly similar cluster genomes. Sample named on the main tree is the one putative cluster sample that was excluded from the cluster based on genome sequence. **B.** Distance matrix showing pairwise differences between the 17 complete genomes assembled from this sample set. Putative cluster samples are bolded. **C.** Spike-in counts for each of the 24 samples and water controls in this sequencing batch.

Analysis of the SDSIs confirmed that genome sequence similarity among cluster-associated samples was not the result of cross-contamination from highly amplified final libraries **(Fig 5c)**. Indeed, 23/24 libraries (22 patient samples and 2 water controls) contained >99% SDSI-mapped reads corresponding to the expected SDSI. One sample (MA_MGH_02845) showed 18% of reads from a second SDSI, which was added to a different sample in the batch (MA_MGH_02839). Upon manually curating the genome from MA_MGH_02845, we first compared it to that of the potentially contaminating sample (MA_MGH_02839). We were not able to assess whether these genomes might share SNVs because there was no genome assembled for MA_MGH_02839; furthermore, we did not identify either sample as cluster-related. Had these genomes shared SNVs or had we identified the suspected contaminated sample as cluster-related, we may have opted to re-sequence. In this way, SDSIs can increase confidence in cluster identification.

## Discussion / Conclusion

As the SARS-CoV-2 pandemic continues and new genomic variants increasingly emerge, it is imperative to build robust experimental confidence into genomic surveillance data interpretation. Here we report the design and implementation of Synthetic DNA Spike-ins (SDSI) as an essential component for tracking and tracing contamination, a potential confounder in amplicon-based SARS-CoV-2 sequencing methods. Our *in silico* design generated robust synthetic targets while mitigating inter-spike-in sequence homology as well as homology with human, SARS-CoV-2, and common laboratory reagents. SDSIs can readily be adopted by laboratories and platforms of all sizes with only minor changes to existing methodologies, little additional cost per sample ($0.006 in our hands), and no interruption or addition of time to standard workflow methodologies. While broadly applicable to most amplicon-based approaches, we coupled the SDSIs to an improved ARTIC amplicon sequencing protocol for recovering genomes from low viral load samples.

Amplicon-based sequencing methods fill a critical need for rapid turnaround and full genome recovery for epidemiological surveillance and clinical applications where SNV identification is crucial. The SDSI+ARTIC protocol which excelled in genome recovery also demonstrated genome concordance with the gold standard approach of unbiased sequencing. Although there was still considerable non-uniformity for samples with low viral loads, small protocol alterations such as changes in the annealing temperature to recover poorly performing amplicon 64 ^7^, primer sequence alteration ^29^,or using 2x primer concentrations for a subset of underperforming amplicons improve overall performance and yield more even coverage. Alternative approaches for the recovery of genomes from samples with low viral load include the use of targeted enrichment approaches ^30,31^ which are more costly and time-consuming. Recent reports of SARS-CoV-2 variants that can affect viral transmissibility, virulence, and susceptibility to pre-existing immunity pose a need for rapid and accurate viral genome sequencing to inform patient care and infection control interventions. Our application of SDSI+ARTIC confirming a cluster representing nosocomial transmission further emphasizes the utility of the SDSIs to confidently identify samples of high genetic similarity.

More broadly, standardizing controls across the viral surveillance community would increase accuracy and integrity of SARS-CoV-2 genomic data worldwide. These SDSIs not only enable profiling of in-batch contamination, but also laboratory-wide detection as their presence in other data (amplicon sequencing, unbiased sequencing, qPCR, or otherwise) would indicate a tagged amplification and thus contamination. Additional synthetic targets could be designed using the same principles to expand into 384 well formats and beyond. Moreover, the approach is applicable to both Illumina and Nanopore sequencing platforms as well as any other existing or future tiled amplicon panel, such as those previously used for Zika, Ebola, and other recent outbreaks ^27,32^. Primer sites could also be easily adapted for integration with new advancements in amplicon sequencing, like tailed primer approaches ^8^. SDSIs could serve as a broad tool for tracing potential contamination across a plethora of fields that employ amplicon based genomic sequencing, such as food safety, species identification and environmental sampling. Altogether, we believe that integration of SDSIs mitigates a critical vulnerability of amplicon-based sequencing, while preserving the many advantages, increasing the robustness of its use across laboratory and clinical settings.

## Acknowledgements

We gratefully acknowledge the microbiology lab staff and infection control personnel at MGH and DPH and all members of the COVID-19 emergency response efforts at MGH, BHCHP, and MADPH. We thank Patricia Rodgers and the entire Broad Flow Core Team for sharing laboratory space and equipment. We also thank Kayla Barnes, Sid Raju, and Sameed Siddiqui for valuable feedback and helpful discussions. Funding: This work was sponsored by the National Institute of Allergy and Infectious Diseases (U19AI110818 to P.C.S.), Centers for Disease Control Broad Agency Announcement (75D30120C09605 to B.L.M), the Bill and Melinda Gates Foundation (Broad Institute), and the US Food and Drug Administration (HHSF223201810172C), with in-kind support from Illumina, Inc., as well as support from the Herchel Smith Fellowship (K.A.L.), the Doris Duke Charitable Foundation (J.E.L.), the Howard Hughes Medical Institute (P.C.S.), and the National Human Genome Research Institute (K99HG010669 to S.K.R.). This work is made possible by support from Flu Lab and a cohort of generous donors through TED’s Audacious Project, including the ELMA Foundation, MacKenzie Scott, the Skoll Foundation, and Open Philanthropy. The content is solely the responsibility of the authors and does not necessarily represent the official views of the National Institutes of Health.

## Author Contributions

G.A., M.R.B., K.A.L, E.N, L.P., S.K.R., A.G.Y. performed laboratory experiments. K.F., C.L., K.A.L., E.N., K.J.S. designed and performed data analysis. J.E.L., D.H., V.P., B.S., E.S. identified and provided samples. S.K.R., K.J.S. conceived of study. M.R.B., K.A.L., E.N., S.K.R., K.J.S. drafted the manuscript. B.M., D.P., K.J.S., P.C.S. secured resources and oversaw study implementation. All authors reviewed and approved the manuscript.

## Methods

We have provided the protocol on Benchling for public use in addition to the detailed methods below: https://benchling.com/s/prt-R95g0tCxKOeCAqn8lAk3

### SDSI design and in silico validation

We designed 96 synthetic DNA fragments that each contained a 140 bp unique sequence and constant priming regions. SDSI homology to sequences from various organisms was predicted by a permissive BLAST search (blastn; 5000 max targets; E=10; word size=11; no mask for low complexity). For different analyses, we modulated filters for percent identity and query cover, and filtered results to various taxa, including *homo sapiens* (taxid:9606), SARS-CoV-2 (taxid:2697049), and archaea (taxid:2157). When we performed the permissive BLAST search described above on the 140 bp core SDSI sequences and filtered results to *homo sapiens* and SARS-CoV-2 with >50 percent identity and >50 query cover, there were no significant hits. When we instead filtered results to exclude archaea with >75 percent identity and >75 query cover, the genus of any significant hit was noted in **Sup Table 1**. We calculated the length and GC content of SDSI primers and amplicons using Geneious Prime (2019.2.1). For every pairwise combination of core SDSI sequences, Hamming distance was calculated by summing the number of mismatched bps between the two 140 bp sequences.

### SDSI application to ARTIC SARS-CoV-2 sequencing

We performed PCR on each SDSI oligo, using the standard SDSI+ARTIC PCR conditions (https://benchling.com/s/prt-R95g0tCxKOeCAqn8lAk3), then ran the PCR products on a 2.2% agarose gel to confirm that these primers amplified the SDSIs and that the product was clean and of the expected size (**Sup Fig 1b**). We also performed PCR with 0.17X SYBR Green added to the mix to perform qPCR assays. We performed this qPCR with multiple templates: (1) 0.5μL of a representative SDSI (1pM), (2) 0.5μL of a representative SDSI + 0.5μL of cDNA from an NP swab, (3) 0.5μL of cDNA from an NP swab, and (4) no template (**Sup Fig 1c**). The oligos were stored at 10μM. To determine an optimal concentration, we tested these SDSIs further diluted to 1, 0.1, 0.01, and 0.001fM; 1μM was added to 5μL of cDNA, to be split to 2×3μL for each ARTIC pool. SDSI primers were added to each ARTIC pool with a final concentration of 40nM. We continued to use primers at this concentration and SDSIs were diluted to 1fM for the full validation batch and cluster investigation.

There are 48 SDSIs that we designed and tested but did not further explored here because they were identified in multiple samples nonspecifically, indicating either that their sequence had homology to other sequences in the reaction, or that there was contamination during oligo synthesis, dilution, or sample processing. We have also ordered a set of synthetic DNA constructs with a T7 polymerase promoter, and may consider transcribing RNA, which could be added to samples earlier in the processing pipeline, including directly into biosample.

### Sample collection and study design

This study was approved by the Partners Institutional Review Board under protocol 2019P003305 and we obtained samples under a waiver of consent for viral sequencing. All samples were excess specimens from clinical testing from the MGH Microbiology Laboratory. All samples were nasopharyngeal swabs in either MTM or VTM. All samples used for method optimization and validation are part of a previously reported cohort^1^. These unique biological materials are not available to other researchers as they are human patient samples from clinical excess material and thus are of limited volume.

### RT, PCR enzyme, and condition optimization

We tested reverse transcriptase enzymes using extracted RNA from SARS-CoV-2 positive clinical samples (CT 13-33). We added 2μL of purified DNase treated RNA as input into SuperScript III (Thermo #18080093), SuperScript IV (Thermo #18091050), or SuperScript IV VILO (Thermo #11756500). Superscript IV reactions incubated at room temperature for 10 minutes, followed by 50°C for 60 minutes and an inactivation step at 80°C for 10min. Superscript IV VILO shared the same protocol, but with a temperature of 85°C for the inactivation step. We input 2.5μL of cDNA for ARTIC pool #1 PCR under standard conditions for 40 cycles. We then tested the resulting pool #1 using the modified Illumina DNA Flex library construction and sequenced on Illumina Miseq (V2 reagent kit) with 2 x 150 bp paired end sequencing.

We tested PCR enzyme efficiency using extracted RNA from SARS-CoV-2 positive clinical samples followed by cDNA generation using SuperScript IV and diluted the resulting cDNA to a mock CT value of 35 for standardization across all PCR enzyme tests. We set up the standard ARTIC PCR pool #1 and pool #2 using an input of 2.5μL, altering only the PCR enzyme and corresponding buffer. We also tested NEB Q5 Hot Start High-fidelity 2x Master Mix (NEB #M0494L), NEB Q5 Hot Start High-fidelity 2x Master Mix plus .01% SDS, NEB Q5 Ultra II Master Mix (NEB #M0544L), KAPA HiFi HotStart (Roche #KK2601), and KOD Hot Start DNA polymerase (Sigma-Aldrich #71842), quantifying resulting ARTIC PCR amplicons using High Sensitivity DNA Qubit, then inputting into modified Illumina DNA Flex library construction. The resulting libraries (except Q5 plus .01% SDS, which had no yield via agarose gel image) were quantified and pooled on Illumina Miseq (V2 reagent kit) with 2 x 150 paired end sequencing.

We optimized PCR cycling conditions on mock CT 35 cDNA (generated as described above) using standard ARTIC PCR primer conditions and .5X primer concentrations. We performed a catch-up/rehybridization PCR under the following conditions: 98°C for 30s, 95°C for 15s then 65°C for 5 min (10 cycles), 95°C for 15s then 80°C for 30s then 65°C for 5 min (2 cycles), 95°C for 15s then 65°C for 5 min (8 cycles), 4°C Hold. We quantified the resulting ARTIC PCR amplicons using High Sensitivity DNA Qubit and input into modified Illumina DNA Flex library construction. We then quantified these libraries and pooled on Illumina Miseq (V2 reagent kit) with 2 x 150 paired end sequencing.

### Illumina library construction comparison

We performed a head-to-head comparison of standard Illumina Nextera DNA Flex and Nextera XT library construction kits. We performed each on post ARTIC v1 PCR amplicons from clinical samples. In short, we amplified samples from a varying range of CT values with ARTIC v1 primers, producing 400 bp size fragments. We then quantified amplicons from each ARTIC primer pool and pooled in equal molar concentrations. Standard Nextera DNA Flex input was 100ng (50ng from each pool) and 1ng (.5ng from each pool) for Nextera XT. We quantified and pooled the resulting libraries before sequencing on an Illumina Miseq (V2 reagent kit) with 2 × 150 paired end sequencing.

We optimized Illumina DNA Flex library construction with the goal of reducing normalization steps and increasing throughput. We scaled down (.5X) Illumina DNA Flex throughout the standard Illumina sequencing protocol, also scaling down sample input for a total of 50ng (25ng from each primer pool). To further reduce steps for normalization, we made a CT dilution before ARTIC PCR standardizing a CT of 27 as input across all samples with a CT of 27 or below. We calculated the difference between 27 and the viral CT then rounded to the nearest whole number. Calculations assumed 100% PCR efficiency. We calculated the number of doublings required for a CT 27, and performed this dilution on all samples pre-ARTIC PCR, removing the pre-DNA Flex DNA concentration and pooling step. We used 1-2μL of post ARTIC PCR amplicon as input into the modified DNA Flex library construction, and performed post library construction quantification and pooling with more uniform library size and concentration, further reducing time and cost of pooling libraries for sequencing.

### Cycle Test

We further optimized ARTIC PCR by modifying PCR cycle numbers. Extracted RNA from SARS-CoV-2 positive clinical samples ranging from CT 27-37 were converted to cDNA with Superscript IV and amplified under standard ARTIC PCR reaction components (with Q5 2x MasterMix) modifying the final number of cycles of PCR from 35, 40 and 45. We quantified cDNA and used at a standard 50ng of input for modified Illumina DNA Flex Library Construction, then quantified the resulting libraries and pooled on Illumina Miseq (V2 reagent kit) with 2 × 150 paired end sequencing.

### Ramp Test

We used Mock CT 35 to test the effect of decreased ramp speed on genome recovery and coverage (**Sup Fig 4c**). Normal ARTIC PCR conditions for this experiment were 98°C for 30 seconds, followed by 40 cycles of 95°C for 15 seconds and 65°C for 5 minutes with a cooling and heating ramping speed of 3°C/s. We tested a slow ramp PCR protocol with the ramp speed reduced to 1.5°C/s. Slower ramp speeds have been shown to reduce GC bias and result in greater coverage^33^. Samples underwent our modified DNA Flex library construction and were sequenced on Illumina Miseq (V2 reagent kit) with 2 × 150 paired end sequencing.

### Primer Concentration Optimization

Under standard ARTIC protocol conditions, we ordered lyophilized ARTIC v3 primers from IDT and resuspended in water at 100μM each. Pool #1 primers consisted of all odd numbered amplicons whereas pool #2 primers consisted of all even numbered amplicons. To generate the 100μM pool #1 primer stock, we combined 5μL of each 100μM pool #1 primer, and repeated this protocol for the even numbered primers to give a 100μM pool #2 primer stock. We selected a total of 20 amplicons as regions of low coverage from previous sequencing data (**Sup Table 3**). Low coverage amplicons were present in both pools, with 11 coming from pool #1 and 9 coming from pool #2. For the primer 2x pools, we spiked in primers for the corresponding amplicons at 2x the concentration (20.8nM final) of the other primers in the pool. For these low coverage primers, we used 10μL of the 100μM stock rather than 5μL. We diluted both the original and 2x primer pools 1:10 in nuclease free water to generate a 10μM working stock. We then selected 8 samples with varying CT values to determine if selectively increasing primer concentrations reduced amplicon dropout. We used our modified ARTIC protocol and processed each sample with both the original primer pool, as well as the 2x primer pool, then sequenced these 16 samples on an Illumina Miseq (V2 reagent kit) with 2 x 150 paired end sequencing.

### Paragon

We used the Paragon CleanPlex SARS-CoV-2 Research and Surveillance Panel protocol to process five RNA samples (CT= 20, 25, 30, 35, and 37)(https://www.paragongenomics.com/wp-content/uploads/2020/03/UG4001-01_-CleanPlex-SARS-CoV-2-Panel-User-Guide.pdf). To make cDNA, we performed reverse transcription using SuperScript IV, and 5μL of this cDNA was used for the multiplex PCR reaction. We amplified samples for 24 cycles per the recommended protocol and pooled in equal concentrations with the corresponding ARTIC samples, then sequenced on a NovaSeq SP and analyzed as described below.

### Metagenomic sequencing and comparison

Metagenomic sequencing data and genome assemblies used for the comparison of amplicon-based sequencing were previously prepared, sequenced, analyzed as described previously,^1^ and the data are publicly available at NCBI’s GenBank and SRA databases under BioProject PRJNA622837. We prepared amplicon sequencing libraries following our SDSI+ARTIC amplicon sequencing protocol (**Fig 1**). Our modified ARTIC pipeline began synthesizing cDNA on 2.5μL of DNAsed RNA using Superscript IV. In order to increase sample throughput and bypass an additional more laborious quantification step post the ARTIC PCR, we normalized cDNA samples that had a high viral load (CT<27) to a CT of 27. To prepare for the ARTIC PCR, we transferred 5μL of the normalized cDNA to a new plate and added 1μL of a SDSI. After mixing, we transferred 3μL to a new plate, added ARTIC PCR pool #1 mastermix and pool #2 mastermix to the respective plates, and on a thermal cycler incubated at 98°C for 30s, followed by 40 cycles of 95°C for 15s and 65°C for 5min. We then combined in equal molar amounts of amplified samples for a total of 50ng and processed through .5X Illumina Flex library construction pipeline. We sequenced the concordance data set on a NovaSeq SP and analyzed as detailed in the methods below.

### Suspected nosocomial cluster investigation

We received NP swab samples in UTM and extracted RNA from 200μL of biosample as previously described^1^. We prepared amplicon sequencing libraries as described above and analyzed as detailed in the methods below. A pairwise distance was calculated between all partial genomes (>80% complete), excluding gaps, to determine whether samples were likely to be the result of nosocomial transmission. We calculated the proportion of reads that mapped to a given SDSI out of all reads that mapped to any SDSI. Metagenomic sequencing libraries, used to confirm genome assemblies, were prepared, sequenced and analyzed as described in^1^. Data has been made available in both the Short Read Archive and NCBI GenBank under Bioproject PRJNA622837. GenBank accessions for SARS-CoV-2 genomes from this set of samples are MW454553 - MW454562.

### Computational analysis workflow

We analyzed sequencing data on the Terra platform (app.terra.bio) using viral-ngs 2.1.1 with workflows that are publicly available on the Dockstore Tool Repository Service (dockstore.org/organizations/BroadInstitute/collections/pgs).

Samples were demultiplexed using the demux_plus workflow with a spike in database file for the SDSIs. We performed any separate analyses to quantify read counts, including those for SDSIs, with the align_and_count_multiple_report workflow with the relevant database. For most analyses involving direct comparisons between samples, we performed downsampling to the lowest number of reads passing filter with the downsample workflow. We performed assembly using the assemble_refbased workflow to the following reference fasta: *https://www.ncbi.nlm.nih.gov/nuccore/NC_045512.2?report=fasta.* We used iVar version 1.2.1 for primer trimming on all samples followed by assembly with minimap2 set to a minimum coverage of either 3, 10, or 20, skipping deduplication procedures. For the Paragon analysis, we trimmed primers using the coordinates of the Paragon primers and assembled samples in the assemble_refbased workflow with novoalign.

### Phylogenetic tree reconstruction

To place the suspected nosocomial cluster in a broader genomic context, we performed a subsampling of the genome sequences available in GISAID (as of January 26 2021). We used the sarscov2_nextstrain workflow to perform a Massachusetts-weighted subsampling of samples from 1 November 2020 - 1 November 2021. Our subsampled dataset included 3146 sequences; 1449 samples from Massachusetts, 1425 samples from elsewhere in the United States and 283 from other countries. We constructed a maximum likelihood tree using iqtree with a GTR substitution model and edited and interpreted the tree in Figtree v1.4.4.

## Code availability

Viral genomes were processed using the Terra platform (app.terra.bio) using viral-ngs 2.1.1 with workflows that are publicly available on the Dockstore Tool Repository Service (dockstore.org/organizations/BroadInstitute/collections/pgs). Downstream analyses were performed using Geneious or standard R packages. Custom scripts used to generate figures are available upon request. Methods

## Data availability

Sequences and genome assembly data are publicly available on NCBI’s Genbank and SRA databases under BioProject PRJNA622837. GenBank accessions for SARS-CoV-2 genomes newly reported in this study are MW454553 - MW454562.

## References

1. Lemieux, J. E. et al. Phylogenetic analysis of SARS-CoV-2 in Boston highlights the impact of superspreading events. Science 371, (2021).

2. Popa, A. et al. Genomic epidemiology of superspreading events in Austria reveals mutational dynamics and transmission properties of SARS-CoV-2. Sci. Transl. Med. 12, (2020).

3. Volz, E. et al. Transmission of SARS-CoV-2 Lineage B.1.1.7 in England: Insights from linking epidemiological and genetic data. bioRxiv (2021) doi:10.1101/2020.12.30.20249034.

4. Washington, N. L. et al. Genomic epidemiology identifies emergence and rapid transmission of SARS-CoV-2 B.1.1.7 in the United States. medRxiv (2021) doi:10.1101/2021.02.06.21251159.

5. Chiara, M. et al. Next generation sequencing of SARS-CoV-2 genomes: challenges, applications and opportunities. Brief. Bioinform. (2020) doi:10.1093/bib/bbaa297.

6. Quick, J. et al. Multiplex PCR method for MinION and Illumina sequencing of Zika and other virus genomes directly from clinical samples. Nat. Protoc. 12, 1261–1276 (2017).

7. Tyson, J. R. et al. Improvements to the ARTIC multiplex PCR method for SARS-CoV-2 genome sequencing using nanopore. bioRxiv (2020) doi:10.1101/2020.09.04.283077.

8. Gohl, D. M. et al. A rapid, cost-effective tailed amplicon method for sequencing SARS-CoV-2. BMC Genomics 21, 863 (2020).

9. Baker, D. J. et al. CoronaHiT: large scale multiplexing of SARS-CoV-2 genomes using Nanopore sequencing. Cold Spring Harbor Laboratory 2020.06.24.162156 (2020) doi:10.1101/2020.06.24.162156.

10. Wu, K. J. These Researchers Tested Positive. But the Virus Wasn’t the Cause. The New York Times (2020).

11. Boers, S. A., Jansen, R. & Hays, J. P. Understanding and overcoming the pitfalls and biases of next-generation sequencing (NGS) methods for use in the routine clinical microbiological diagnostic laboratory. European Journal of Clinical Microbiology & Infectious Diseases vol. 38 1059–1070 (2019).

12. Charre, C. et al. Evaluation of NGS-based approaches for SARS-CoV-2 whole genome characterisation. Virus Evol 6, veaa075 (2020).

13. Rausch, J. W., Capoferri, A. A., Katusiime, M. G., Patro, S. C. & Kearney, M. F. Low genetic diversity may be an Achilles heel of SARS-CoV-2. Proceedings of the National Academy of Sciences of the United States of America vol. 117 24614–24616 (2020).

14. Endo, A., Centre for the Mathematical Modelling of Infectious Diseases COVID-19 Working Group, Abbott, S., Kucharski, A. J. & Funk, S. Estimating the overdispersion in COVID-19 transmission using outbreak sizes outside China. Wellcome Open Res 5, 67 (2020).

15. Wong, F. & Collins, J. J. Evidence that coronavirus superspreading is fat-tailed. Proceedings of the National Academy of Sciences vol. 117 29416–29418 (2020).

16. Dearlove, B. et al. A SARS-CoV-2 vaccine candidate would likely match all currently circulating variants. Proceedings of the National Academy of Sciences vol. 117 23652–23662 (2020).

17. Adam, D. C. et al. Clustering and superspreading potential of SARS-CoV-2 infections in Hong Kong. Nature Medicine vol. 26 1714–1719 (2020).

18. Organization, W. H. & Others. Genomic sequencing of SARS-CoV-2: a guide to implementation for maximum impact on public health, 8 January 2021. (2021).

19. COVID-19 Genomics UK (COG-UK). An integrated national scale SARS-CoV-2 genomic surveillance network. Lancet Microbe 1, e99–e100 (2020).

20. Lagerborg, K. A., Watrous, J. D., Cheng, S. & Jain, M. High-Throughput Measure of Bioactive Lipids Using Non-targeted Mass Spectrometry. Methods Mol. Biol. 1862, 17–35 (2019).

21. Boja, E. S. & Rodriguez, H. Mass spectrometry-based targeted quantitative proteomics: achieving sensitive and reproducible detection of proteins. Proteomics 12, 1093–1110 (2012).

22. Chen, K. et al. The Overlooked Fact: Fundamental Need for Spike-In Control for Virtually All Genome-Wide Analyses. Molecular and Cellular Biology vol. 36 662–667 (2016).

23. Illumina COVIDSeq Test. https://emea.illumina.com/products/by-type/ivd-products/covidseq.html

24. Dilucca, M., Forcelloni, S., Pavlopoulou, A., Georgakilas, A. G. & Giansanti, A. Codon usage and evolutionary rates of the 2019-nCoV genes. Cold Spring Harbor Laboratory 2020.03.25.006569 (2020) doi:10.1101/2020.03.25.006569.

25. Mathieu-Daudé, F., Welsh, J., Vogt, T. & McClelland, M. DNA Rehybridization During PCR: The ‘C o t Effect’ and Its Consequences. Nucleic Acids Res. 24, 2080–2086 (1996).

26. Klempt, P. et al. Performance of Targeted Library Preparation Solutions for SARS-CoV-2 Whole Genome Analysis. Diagnostics (Basel) 10, (2020).

27. Metsky, H. C. et al. Zika virus evolution and spread in the Americas. Nature 546, 411–415 (2017).

28. So, A. P. et al. A robust targeted sequencing approach for low input and variable quality DNA from clinical samples. NPJ Genom Med 3, 2 (2018).

29. Itokawa, K., Sekizuka, T., Hashino, M., Tanaka, R. & Kuroda, M. Disentangling primer interactions improves SARS-CoV-2 genome sequencing by multiplex tiling PCR. PLoS One 15, e0239403 (2020).

30. Metsky, H. C. et al. Capturing sequence diversity in metagenomes with comprehensive and scalable probe design. Nat. Biotechnol. 37, 160–168 (2019).

31. Houldcroft, C. J., Beale, M. A. & Breuer, J. Clinical and biological insights from viral genome sequencing. Nat. Rev. Microbiol. 15, 183–192 (2017).

32. Quick, J. et al. Real-time, portable genome sequencing for Ebola surveillance. Nature 530, 228–232 (2016).

33. Aird, D. et al. Analyzing and minimizing PCR amplification bias in Illumina sequencing libraries. Genome Biol. 12, R18 (2011).

