## Supplemental Materials for "DNA spike-ins enable confident interpretation of SARS-CoV-2 genomic data from amplicon-based sequencing"

### *Supplementary Tables*

**Supplementary Table 1:** Spike-In Design
**Supplementary Table 2:** SDSI and Viral Read Percentages
**Supplementary Table 3:** ARTIC v3 Primers and Primers Spiked in at 2X
**Supplementary Table 4:** Time and Cost Comparison of FLEX vs XT
**Supplementary Table 5:** Cost of SDSI+ARTIC

### *Supplementary Figures*

**Supplemental Figure 1.** Spike-in validation.
**Supplemental Figure 2.** SDSI Titration.
**Supplemental Figure 3.** Comparison to alternate amplicon sequencing strategies.
**Supplemental Figure 4.** Reverse transcription and amplification variations to the ARTIC protocol.
**Supplemental Figure 5.** Increasing primer concentration 2-fold in regions of low amplicon coverage.
**Supplemental Figure 6.** Modified Flex outperforms XT in coverage depth and evenness at lower cost.
**Supplemental Figure 7.** SDSI+ARTIC over a diverse set of samples has superior genome recovery and more coverage uniformity at higher CTs.

### *Supplementary Figures*

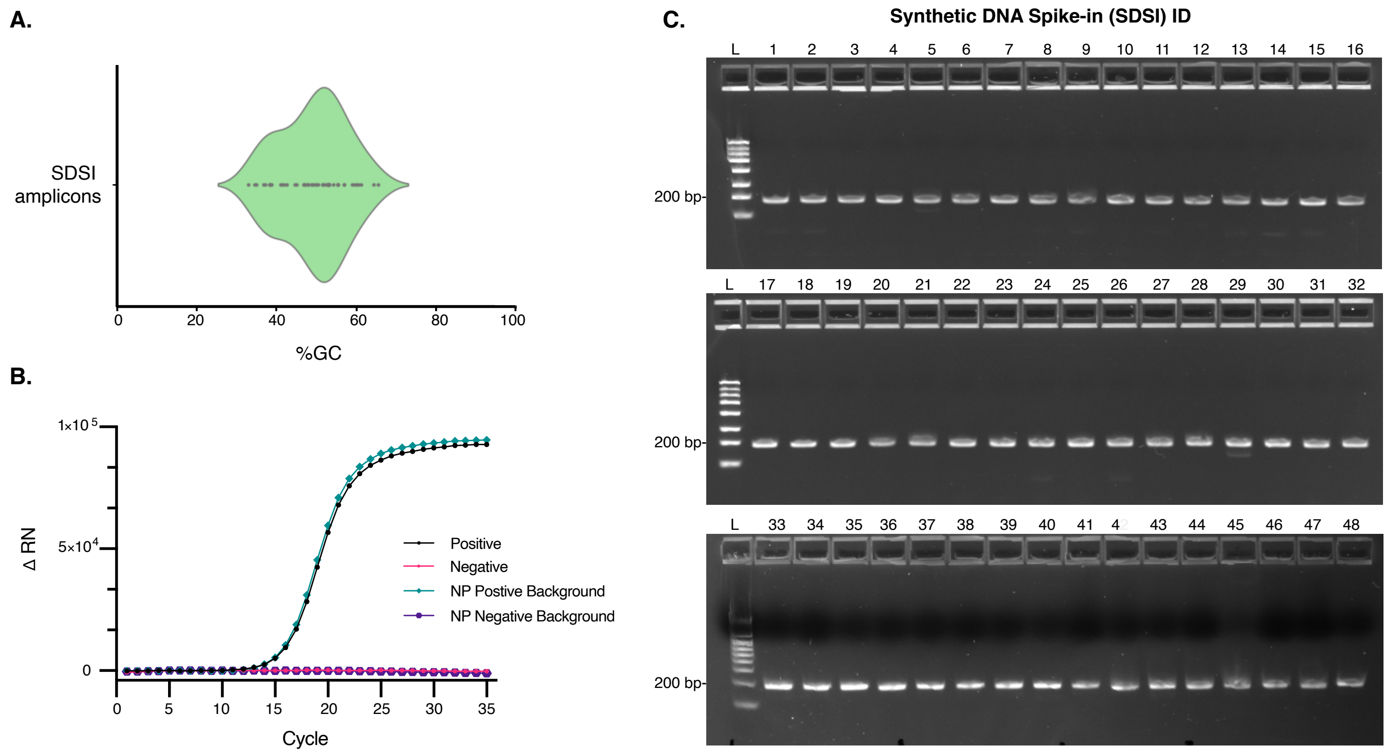

**Sup Figure 1. Spike-in validation.**
**A.** The distribution of GC content of SDSI amplicons. **B.** 100fmol DNA spike-in amplified under standard ARTIC PCR conditions for 40 cycles run on 2.2% agarose gel image with 188bp amplified spike-in (SDSI 1-48) **C.** RT-PCR for Spike-in and spike-in specific primers, Spike-in specific primers water control, Spike-in with COVID positive cDNA and spike-in specific primers, COVID positive cDNA and spike-in specific primer

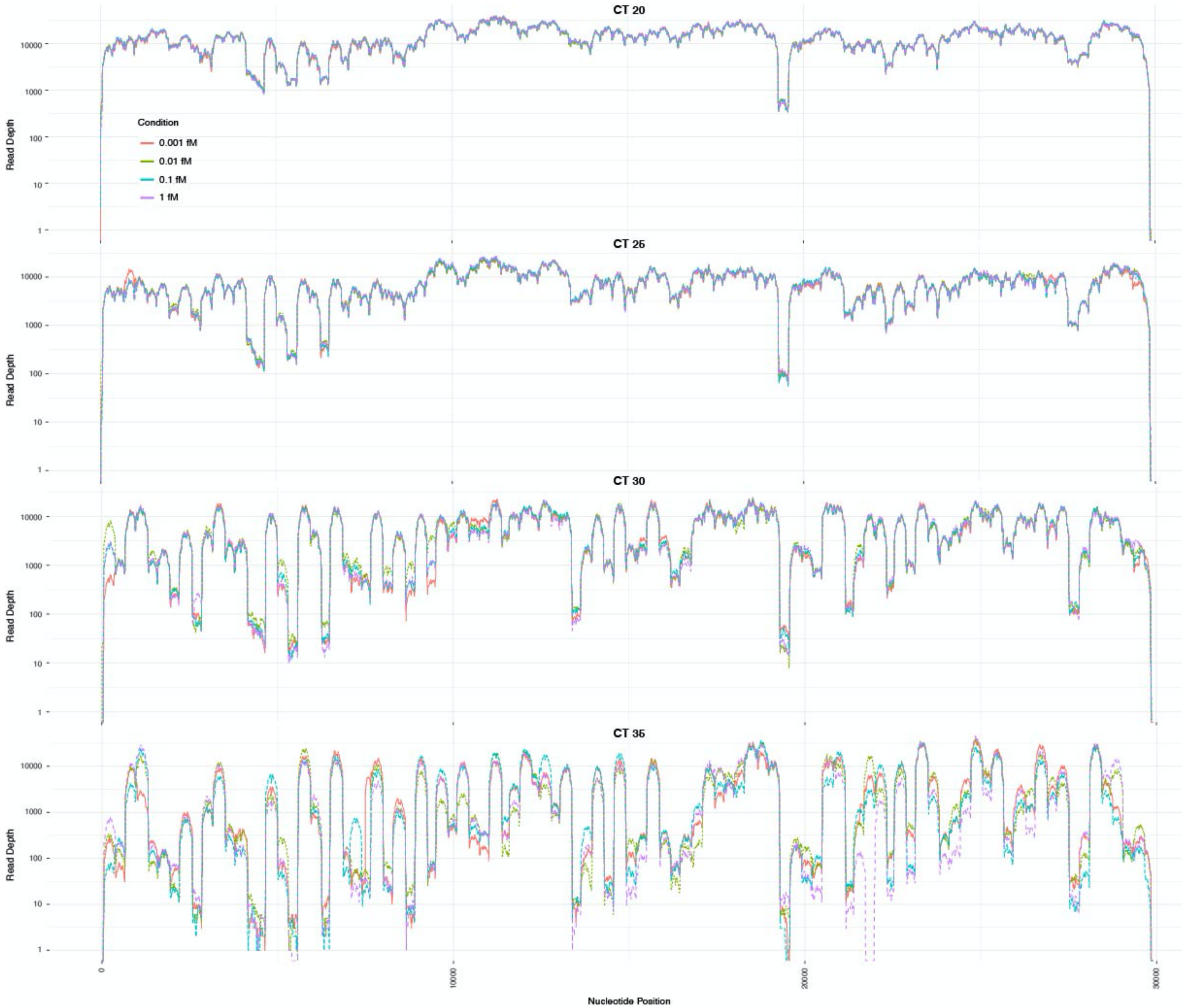

**Sup Figure 2. SDSI Titration.**

Coverage plots for four different SDSI concentrations (1fM, 0.1fM, 0.01fM, 0.001fM) at four different CT dilutions (CT=20,25,30,35).

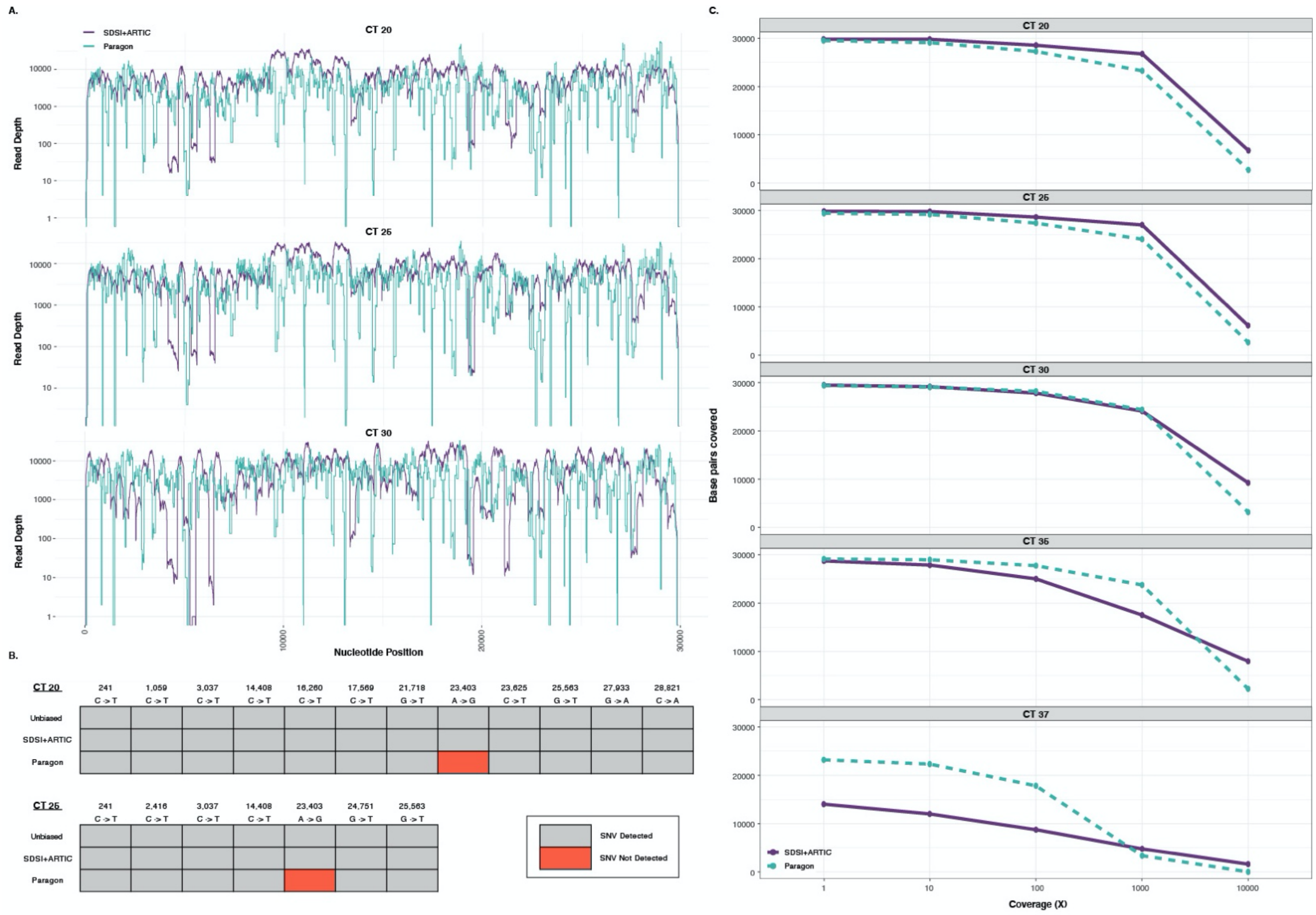

**Sup Figure 3. Comparison to alternate amplicon sequencing strategies.**

**A.** Three representative coverage plots for CT 20, CT 25, and CT 30 samples. **B.** SNV detection for the CT 20 and CT 25 sample. ARTIC and Paragon consensus sequences were compared to our unbiased metagenomic consensus sequences. The SNV that was not called in Paragon was due to low coverage at that position. Analysis was performed with assemblies generated with a minimum coverage of both 3 and 20, yielding identical results. **C.** Base pairs of the SARS-CoV-2 genome covered for our SDSI+ARTIC pipeline versus Paragon CleanPlex Panel at different depths of coverage. Five samples at varying CTs were compared.

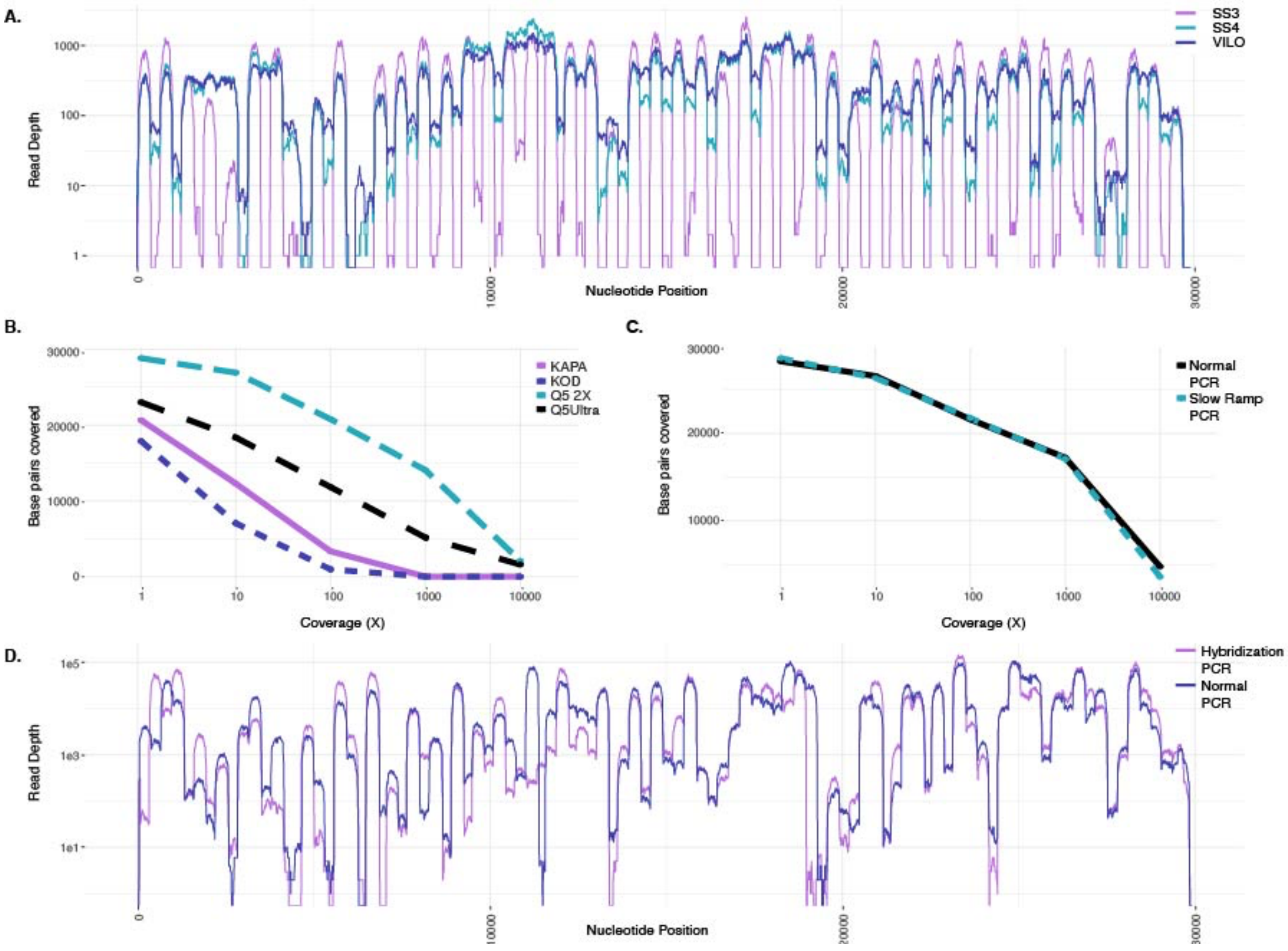

**Sup Figure 4. Reverse transcription and amplification variations to the ARTIC protocol.**

**A.** Read depth across each nucleotides position for the same sample (CT=13.89) when using three different reverse transcriptases (SSIII, SSIV, or SSVILO) for cDNA synthesis. **B.** Base pairs of the SARS-CoV-2 genome covered at various depths when using different enzymes for the ARTIC PCR. **C.** Base pairs of the SARS-CoV-2 genome covered at various depths when using either normal ramping speed (3°C/s) for the ARTIC PCR or reduce the ramping (1.5°C/s). **D.** Read depth across each nucleotides position for normal ARTIC PCR vs an alternate hybridization PCR.

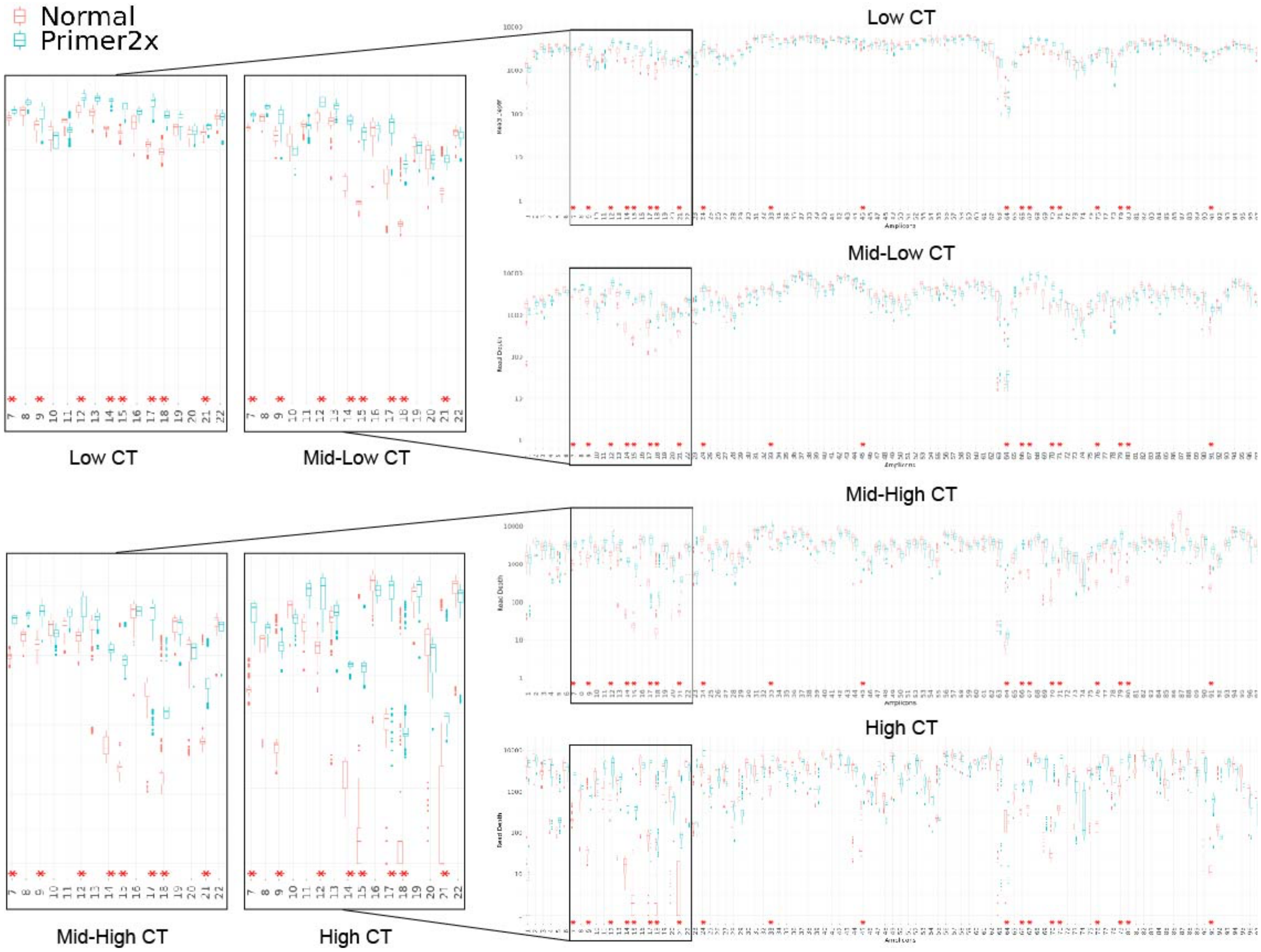

**Sup Figure 5. Increasing primer concentration 2-fold in regions of low amplicon coverage.**
Red asterisk indicates amplicons in which the primer pairs were spiked in at 2X the concentration of the others in the pool. Box plots showing the distribution of absolute sequencing coverage (log10) per amplicon for ARTIC PCR conditions (Normal) and Primer 2x concentrations for 4 representative samples. The boxes are plotted by the Q1, median, and Q3, the whiskers by Q1/Q4, and the outliers by the dots.

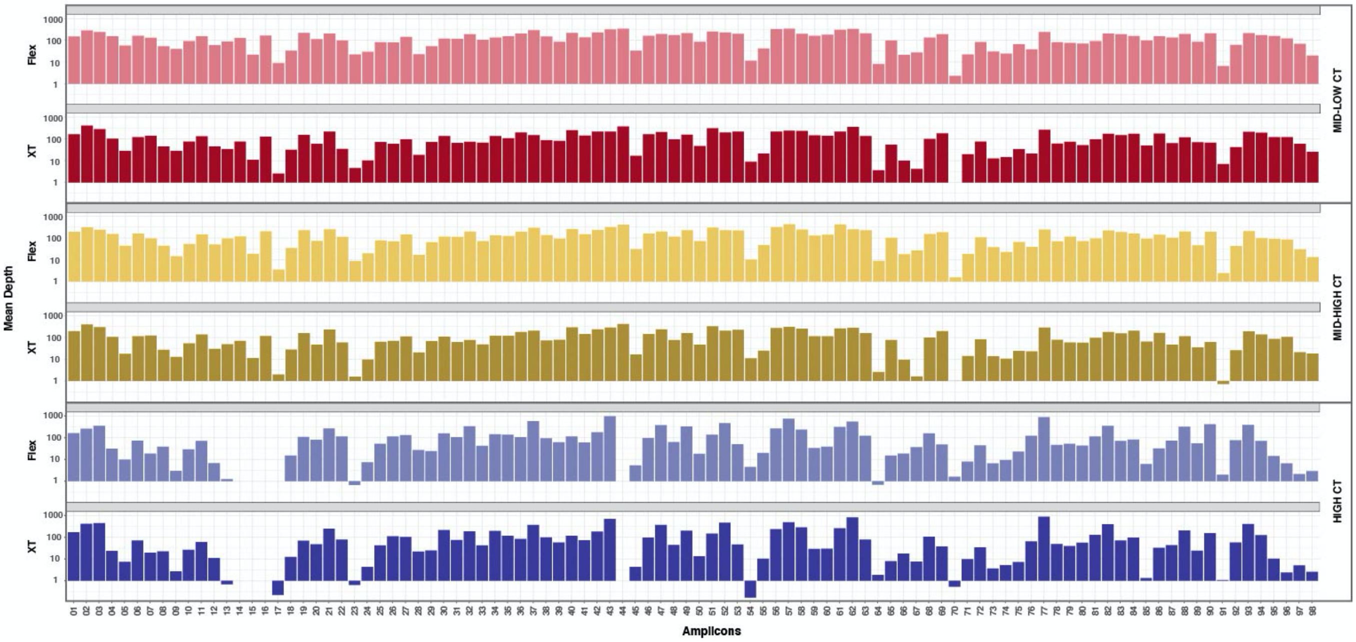

**Sup Figure 6. Modified Flex outperforms XT in coverage depth and evenness at lower cost.**
Illumina Nextera XT and modified Illumina Nextera Flex library construction on three samples with varying CTs. Plotted is the mean sequencing depth (log10) per amplicon.

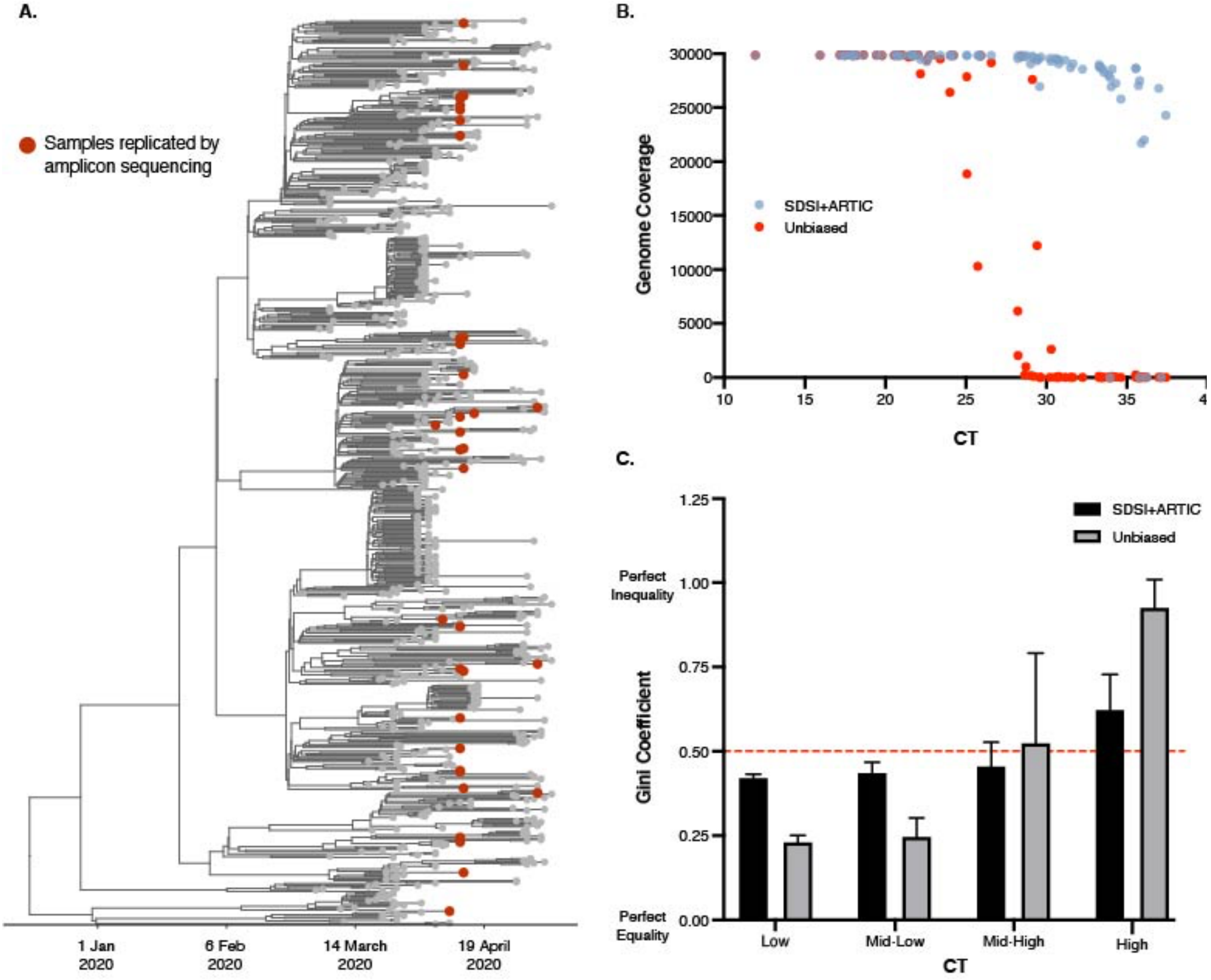

**Sup Figure 7. SDSI+ARTIC over a diverse set of samples has superior genome recovery and more coverage uniformity at higher CTs.**
**A.** Time-measured maximum clade credibility tree of 772 genomes from Massachusetts, reported in Lemieux et al., 2020. The 89 samples compared for metagenomic and amplicon sequencing are shown with red dots. **B.** Genome coverage for unbiased metagenomic sequencing versus SDSI+ARTIC amplicon sequencing pipeline (N=81, excluded samples had no detectable CT). All samples downsampled to 975,000 reads. **C.** Gini coefficients grouped by CT (N=70, excluded samples that did not generate assemblies in either one or both methods). Dashed red line represents the median.

*Supplementary Tables*

### Sup Table 1. Spike-In Design

| Oligo ID | Sequence to order (5' to 3') | Genera with significant homology* (excluding genera from the domain archaea) |
|---|---|---|
| SDSI forward primer | TCTCCTTCTTAGCTTCGTGAGAAC | n/a |
| SDSI reverse primer | CTTGGTCGTCTACTACATGATGTG | n/a |
| SDSI 1 | ACAGTTCTCCTTCTTAGCTTCGTGAGAACGACCGGACGTTGTGATCACGGGTACCTT<br>GATCTGGTACTCAAAGGTTTGCCCCCGTGAAGTCTGGTACATGGCTAGACACGTCAC<br>TCCATTCGAGGGACATTCGAAGTTAGAGAAGGGCAGAGCGATACATCAGATATATCA<br>CATCATGTAGTAGACGACCAAGACAGT | none |
| SDSI 2 | ACAGTTCTCCTTCTTAGCTTCGTGAGAACCTTAATGGAAAGTATGCTTTAGATACCTT<br>CTGGAACGCTATCTCACTTGGCGGGAATTCAGATATGGAGAGTAAATTAAGGGATCT<br>GGAAGTAAAGTTAATGTCGTTAATCTATTTAAATGAGTCACCATTAAAATCACCCACAT<br>CATGTAGTAGACGACCAAGACAGT | none |
| SDSI 3 | ACAGTTCTCCTTCTTAGCTTCGTGAGAACCATAATATGTTAGAGGTAGAATTTCTTTGT<br>GATAGAATATTATTGATGAATGATGGAAGAGAATTAGCATTAGGAAAACCTAAGGAAC<br>TGGTAAAGGATACAGAATCTAAGAATCTTGAAGAGGTTTTCCTTAAACTTGTCACATC<br>ATGTAGTAGACGACCAAGACAGT | none |
| SDSI 4 | ACAGTTCTCCTTCTTAGCTTCGTGAGAACAGTCTAGGTTTTAATTCTTCAACTGCTTCA<br>AATACTAGCTTACTGTAGTTATCTGCCCTCATGTTAGGATATATATCTGGAATATAAGG<br>AGGTTGATGAGTTATAAGAAGTGGATGAAATTGTTGTCACACACTCCCCTACACATCA<br>TGTAGTAGACGACCAAGACAGT | none |
| SDSI 5 | ACAGTTCTCCTTCTTAGCTTCGTGAGAACCTCGTAAGCGTTTCCTACCCTCGAGAGG<br>GCCATCCTGGTGGTGAGGAAGTCGTCGAAGTGGGCTAAGTAAAAAGCGAAGATCTC<br>GACCCACAATTACCTCCTCCTGTACACCAGGAATACCCCTATCAGGATAGAGATACC<br>ACATCATGTAGTAGACGACCAAGACAGT | none |
| SDSI 6 | ACAGTTCTCCTTCTTAGCTTCGTGAGAACTCACGGTCCGCGACGTGAATCGGGCGTT<br>CCAGTCGGCGTTCGGCTACGACGCCGACGACGTGGTCGGAAGCGACCTCCTCGGG<br>CGAATCGTGCCCCCGGTGCCGGACCCGGACCCGGTGCCGGAACCGGGGGACGAC<br>GAGCACATCATGTAGTAGACGACCAAGACAGT | *Nocardioides; Rathayibacter* |
| SDSI 7 | ACAGTTCTCCTTCTTAGCTTCGTGAGAACGCGTCCGCGAGTTCATCCTGAACGTCGT<br>CCCGCTGTCGCCCGGCGAGGAGCGCGGGGCGGGCTACGCCATCTACACCGACATC<br>ACGGAGCGGAAGACCCGCGAAAGCGAGCTAGAGCGACAGAACGAGCGATTGGAGG<br>AGCACATCATGTAGTAGACGACCAAGACAGT | none |
| SDSI 8 | ACAGTTCTCCTTCTTAGCTTCGTGAGAACACGAACTCGTCGGTGAACATCTCGTCTTC<br>CGGGGAGCCCGCCGCTCATGGCCTGCCCCCGCCGTAAGCTGCTGCATAAACCCGC<br>TCCAAAATATACGGATCATTCACCCCTTGGAATCGCTCAATCAGATCAATGTACACCA<br>CATCATGTAGTAGACGACCAAGACAGT | none |
| SDSI 9 | ACAGTTCTCCTTCTTAGCTTCGTGAGAACTGCGTACATTCCCCCTAAGCGGCTCCCA<br>ATATACAGACGCCGGTTAACGACAGCTGGCGACCCTGTGATCTCAGTACCGGTGTC<br>GAATGACCACATCAGCTTGCCTGTCCGTGCATGGAGTTCGTATACGTACCCGTCGTC<br>ACATCATGTAGTAGACGACCAAGACAGT | none |
| SDSI 10 | ACAGTTCTCCTTCTTAGCTTCGTGAGAACATACACCACCCCATCAGCAACAACTGAAT<br>CATGATTAAGTATCGCACCAGCATCGTAGCGCCAGCGTTCACTGCCAGTGGTGCTAT<br>CGAATGCATAGAAGATATGCTCCTAATCGCCAATATCAGTACTTCACAAAGCCGCACA<br>TCATGTAGTAGACGACCAAGACAGT | none |
| SDSI 11 | ACAGTTCTCCTTCTTAGCTTCGTGAGAACGTGGAGTCTTTTGTCACACCGCAGAGGC<br>GTAGCGCTGCAGAGCAGGAGCCCAAGCCTACTGCCAACATAGAGAACATAGTGGCT<br>ACAGTATCCCTCGACCAGACTCTAGACCTGAACCTCATAGAGAGGAGCATACTGACC<br>ACATCATGTAGTAGACGACCAAGACAGT | none |

| | | |
|---|---|---|
| SDSI 12 | ACAGTTCTCCTTCTTAGCTTCGTGAGAACCGTCGCCTGGGTTAAGAGGATGTTCGGC<br>CTCTCCAAGGCGGGTCACGGAGGCACGCTGGACCCGAAGGTCACCGGCGTCCTCC<br>CCGTAGCCCTGGAGGAAGCAACCAAGGTCATAGGCCTGGTGGTGCACACGAGCAAG<br>GCACATCATGTAGTAGACGACCAAGACAGT | none |
| SDSI 13 | ACAGTTCTCCTTCTTAGCTTCGTGAGAACCGTGGGCGAGATCTACCAGAGGCCGCC<br>GCTCCGCAGCAGTGTTAAGAGAAGCCTCCGCGTCAAGAGGATATACGAGATAGAGC<br>TGCTGGAGTACAACGGCAGGTACGCGCTCATGAGGGTGCTCTGCGAGGCCGGCAC<br>ATCACATCATGTAGTAGACGACCAAGACAGT | none |
| SDSI 14 | ACAGTTCTCCTTCTTAGCTTCGTGAGAACCGCTGGAAGAACGAGGGCAAGGAGGAC<br>CTGCTGCGGAGCTACATCAAGCCCGTCGAGTACGCCGTGAGCCACCTGCCCAAGAT<br>AGTTATACGCGATACCGCGGTGGACGCCATAGCCCATGGCGCGAACCTCGCGGTGC<br>CCACATCATGTAGTAGACGACCAAGACAGT | none |
| SDSI 15 | ACAGTTCTCCTTCTTAGCTTCGTGAGAACGGGAGACCCCAAGGTGACCGGCGTCCTA<br>CCAGTGGGGCTCGCCAACAGCACCAAGGTCATTGGTAATGTTATACATAGTGTTAAA<br>GAATACGTGATGGTTATACAGCTCCACGGCGATGTAGCCGAGCAGGATTTAAGAACA<br>CATCATGTAGTAGACGACCAAGACAGT | none |
| SDSI 16 | ACAGTTCTCCTTCTTAGCTTCGTGAGAACTAGAGGGAAAGACTGTAGCTTTCATTCCT<br>AGGCACGGAAAGAGACACAGAATACCTCCACATAAGATAAATTATAGAGCTAATATAT<br>GGGCATTAAAAGAACTAGGAGTGAAATGGGTCATCTCAGTTTCTGCCGTAGGACACA<br>TCATGTAGTAGACGACCAAGACAGT | none |
| SDSI 17 | ACAGTTCTCCTTCTTAGCTTCGTGAGAACTGAGGGAGCTCAGGAGGACTCGCACGG<br>GGCCCTACAGGGAGGATGAGACACTTGTAAGGCTCCAGGACGTCAGCGAGGCCCT<br>GCTCCTGTGGAGGAGCAACGGGGATGAGAGGTATCTTAGACGCATCGTGCTACCCG<br>TTCACATCATGTAGTAGACGACCAAGACAGT | none |
| SDSI 18 | ACAGTTCTCCTTCTTAGCTTCGTGAGAACGAAACATCTATCGCCCACCTCCCGAAGAT<br>AATGATCTTGGATACAGCTGTCGACGCCATAGCACATGGTGCCAACCTGGCTGCCCC<br>AGGCGTCGCCAGGTTAACCAGGAACATCGCGAAGGGTAGTACCGTAGCGATCCTCA<br>CATCATGTAGTAGACGACCAAGACAGT | none |
| SDSI 19 | ACAGTTCTCCTTCTTAGCTTCGTGAGAACTCGCTATCCCCGTGTACAGCATGGTGGG<br>GGTGCCGATGCCCGGGTAGAACTTGGTGACGCTCTCCAGCTTCTCGAGGACGGTTT<br>CCTTGGGGAGGCTCGCGGTGTCCACGAGGGTTATCGCGTCCTCGGCGCCGTCGCC<br>GCACATCATGTAGTAGACGACCAAGACAGT | none |
| SDSI 20 | ACAGTTCTCCTTCTTAGCTTCGTGAGAACCGAGGACGCGAAGAGCGCGGTGGATGT<br>GGACGCGCCGCCGCACACGTAGCCGTCGAGGTAGCGCGGAACCATCGGCGACATC<br>AGCCCCACGACGCGACCCGAGGCGTTGCCGAGGATCACGTCGAGCGTCACGCGCG<br>GCACACATCATGTAGTAGACGACCAAGACAGT | none |
| SDSI 21 | ACAGTTCTCCTTCTTAGCTTCGTGAGAACTCTATGGTGTAGAACGGGTCGTTGCGGA<br>GCCAGCCTGGCGGCACGTACCGGTCGTCCGCTATCGCCAGCGATCTCTCGAAGAG<br>GTCGAGGTAGGCGGACGCGTTGGCGAACGCCCCGTGTATCACGACGTCTATCCCGC<br>CCACATCATGTAGTAGACGACCAAGACAGT | none |
| SDSI 22 | ACAGTTCTCCTTCTTAGCTTCGTGAGAACCCTACGCCGGGTGCGTAGGAGGGCTCG<br>AGTACATCCATGTCTATACTGATGTATGTTTTACCCAGGTCGCCTAGTGCCAGGGGT<br>CCCTTTAACGCTTCCAGGATAGAGTACACGGTGACGTCTCTAGTCTTCTTCAAGAACA<br>CATCATGTAGTAGACGACCAAGACAGT | none |
| SDSI 23 | ACAGTTCTCCTTCTTAGCTTCGTGAGAACCTACTAGCGTGTCAACGGAGCTCTTCAAC<br>GCCTTTACTATTGGATAGGTTATAAGGTGCTCGCCTCCGAGGAATCCCAGGAGCATG<br>CCGGGATACTCGTCTACAACGCCTTTCACCACGTCACCTATGATTCTTAAAGAGCAC<br>ATCATGTAGTAGACGACCAAGACAGT | none |
| SDSI 24 | ACAGTTCTCCTTCTTAGCTTCGTGAGAACCATAGGTGACATGGGGTTTCCCATTGACT<br>CTATAAAGCCGTATCCTTTAAGCGGAGTGCAATTGGTCTACGCTTTGCTTAACAACAG<br>GTATTTCCTACCGGGTAGAGAGGGCTCGCTCATAGCTTTAGGTAGCGTGACGGCAC<br>ATCATGTAGTAGACGACCAAGACAGT | none |
| SDSI 25 | ACAGTTCTCCTTCTTAGCTTCGTGAGAACGGTATCTCACCGCTTGTCACCATAGTATC<br>CCTCAGGTACTCCAGTATTCTTGAGAGAAACGCACCTAAGCCGGATCTCAGGTTTGA<br>ATCCATAAGAACTATGAGTGAAGCGGGATTGAAGCCCCTGCTGTTTCTAAGACCCAC<br>ATCATGTAGTAGACGACCAAGACAGT | none |
| SDSI 26 | ACAGTTCTCCTTCTTAGCTTCGTGAGAACTAAGGGAGATAGAGAAACGCATCAAAATA<br>CCCTTGGGGAAACTGCGTGCAGGGGTTCAATATGGAGTAGAGGTCTCAGACATAAA<br>GGAGAAGATAGCTGCTTACGCTAGGAGGAAGGGGCTTAAATACTTCCCATCGGCAC<br>ACATCATGTAGTAGACGACCAAGACAGT | none |

| | | |
|---|---|---|
| SDSI 27 | ACAGTTCTCCTTCTTAGCTTCGTGAGAACTGTGAACCTCGTGCCCGGCTCTAAGTCGTGAGGGCTTGCAACATAGGTGGGGAGGAACCCGAGCAACGGGTAAGAAGACAGGATAAGCGGTATCGCTATGAAGAGGGCTGAGAAAAGGACATATACTCCTGAGCCCGTCCCACATCATGTAGTAGACGACCAAGACAGT | none |
| SDSI 28 | ACAGTTCTCCTTCTTAGCTTCGTGAGAACCGAACATGCCTTCCCCGTCTATATAGACCCAGTAGAGTTTAAAAACTTAACCAGAGACGGCTTGTGAGCCGGATCTCTCCCCCGCTAGGCCCTGGATTGGGCTCGCTCCTCCTGGGACCCCGGCCTCCACATGCTCGGGACACATCATGTAGTAGACGACCAAGACAGT | none |
| SDSI 29 | ACAGTTCTCCTTCTTAGCTTCGTGAGAACTCTCGGTTCGGCAATAAGTAATACCAACGAGGTATTACCATGCGCGTGACCAGCAAAGGCCAAGTGACGATCCCAAAGGAGATACGGGATCATTTGGGGATTGGGCCGGGCTCCGAGGTGGAGTTCGTGCCCACAGACGACACATCATGTAGTAGACGACCAAGACAGT | *Mesorhizobium; Rhizobium; Neorhizobium; Aminobacter; Sinorhizobium; Shinella;* |
| SDSI 30 | ACAGTTCTCCTTCTTAGCTTCGTGAGAACCTCGATCATATGGCCGGCACGTTGGACTTGGGAGGCATGACAACGGACGAGTATATGGAGTGGCTGAGGGGTCCACGTGAAGATCTCGACATTGATTGACACAAATGTCCTGATCGATGTTTGGGGTCCTGCCGGACAGGCACATCATGTAGTAGACGACCAAGACAGT | *Mesorhizobium; Neorhizobium* |
| SDSI 31 | ACAGTTCTCCTTCTTAGCTTCGTGAGAACCAGGTGTATTTTACACACCTGGACAGCCAGCATATGATGCTAGCACTCGGTGTCCCCTTATCACGGTTTCCCGCATTGTAAAGTTTTCGCGCCTGCTGCGCCCCGTAGGGCCTGGATTCATGTCTCAGAATCCATCTCCGCACATCATGTAGTAGACGACCAAGACAGT | ‘uncultured bacteria’; ‘uncultured prokaryote’; ‘uncultured microorganism’ |
| SDSI 32 | ACAGTTCTCCTTCTTAGCTTCGTGAGAACCGTAGCCCGCACCTTCCTCTGGTTTAGCACCAGCGGTCCCCACAGAGTACCCATCATCCCGAAGGATATGCTGGCAACAGTGGGCACGGGTCTCGCTCGTTGCCTGACTTAACAGGATGCTTCACAGTACGAACTGACGACACATCATGTAGTAGACGACCAAGACAGT | ‘uncultured bacteria’; ‘uncultured prokaryote’; ‘uncultured organism’ |
| SDSI 33 | ACAGTTCTCCTTCTTAGCTTCGTGAGAACGAAACTTACCTTATCAGTGTCATTAAGCATATTGCTTCCAAGACCCATTGAAGCACTTACATCGTTGATACACAGGTGCCAGGAATAGTATTCCTCAGTCTCACTATAATCCTCGTTGGTGTAGCCTTCAAGAGAGTCAACACATCATGTAGTAGACGACCAAGACAGT | none |
| SDSI 34 | ACAGTTCTCCTTCTTAGCTTCGTGAGAACGTTTAAGCAATTCTTCGGATGAAAGATGGCGCTCTATAGGAATTTGTTCTGGTCTAGCCATAAGGCATTATTTGTACTTAATTAGTAATAAATGTTTAGTTAATGACTATAAATCTGCAATTGGAGTCTCAAATTTTCAACACATCATGTAGTAGACGACCAAGACAGT | none |
| SDSI 35 | ACAGTTCTCCTTCTTAGCTTCGTGAGAACAACATGAAGGATGTGTGTAAGAGGAAACGTTATTAACAGACGTAATCAGGAGGATAGTTATGCCCTAAAAACAGCAGAGTTAAGGTTTAAAAATAAGATAAGAACTCAGTTGAGGTTTATCCATTAATCCCATTAATCCTCACATCATGTAGTAGACGACCAAGACAGT | none |
| SDSI 36 | ACAGTTCTCCTTCTTAGCTTCGTGAGAACGTATCCGCTGATATATCCTGGGGATATAGATCGCTCTGAAATGGTTACATCTATCGGTTTTAAGGACAGTTCCAACACTATTGGACCTTGCAGCTATGACAGGAATAATCTGTTTATCGAGCACAGTTGAATTTGACCTACACATCATGTAGTAGACGACCAAGACAGT | none |
| SDSI 37 | ACAGTTCTCCTTCTTAGCTTCGTGAGAACATATTCCGTATTTCTTATCAAACCGATCGTGAAGATTTGACAAAGGCTTAACTTTAGGGCTCCACTTCTCATTATTAGCCTTAGAATATAAAGCGTAACCGTAAGCCTGAGGAACGTAAAGCTTAGGAGATTCAATCCCGCACATCATGTAGTAGACGACCAAGACAGT | none |
| SDSI 38 | ACAGTTCTCCTTCTTAGCTTCGTGAGAACTAAAATTAGCCGAAGGCTTCCCATTACCGAAAAAGTCGTTTATTAGCTCTTCATCCTTCTTCTCCACGTCCGCCCATTCCTCTCCTTCCCTTGGAATTTTAAGCTCGTCCCAGCTGACTCTTATGGGCAATTCAATATCCCACATCATGTAGTAGACGACCAAGACAGT | none |
| SDSI 39 | ACAGTTCTCCTTCTTAGCTTCGTGAGAACTCCGGAGGAATCTATCATATTAAACCTCCTCAAAATCGCCTCCTCTTGATTGCTTAAAGGCTGTGAATTACAAAGCTTATTTAATGCGTCCCAAAGCGTTAAGTAATAATTATTTATATTAAACACTACTATTTCAGTAGCACATCATGTAGTAGACGACCAAGACAGT | none |
| SDSI 40 | ACAGTTCTCCTTCTTAGCTTCGTGAGAACGTTCCTCCTCAATTCAATTGGACTGAAGGAGGGTACGTTCTGGAAAACAGAGCGTAAAAGAGATATAGAACGTAGTATACACATAGCTGGAAAAAGAACAATCATTAAGACAATAAAGAACTTTATGGAAAAGAGTAGAACACATCATGTAGTAGACGACCAAGACAGT | none |
| SDSI 41 | ACAGTTCTCCTTCTTAGCTTCGTGAGAACTCGTGTAAAGGTTGTATAATTCAAGCCTCAGAACATTTCGAACTCCTTACAAAATCGTTTAAACTTTCTAAGGCATAAATTTACTAGAAATTGTCATTTATGAGAATGTAACTATATAGATGGTAAAATTATTAATCCTCCACATCATGTAGTAGACGACCAAGACAGT | none |

| | | |
|---|---|---|
| SDSI 42 | ACAGTTCTCCTTCTTAGCTTCGTGAGAACGGCTGAAAAATAGGTTCGATCCGCCTCC<br>TCACTTCTTCTCCTTCTTGCCCTCGGCCTCGGAGGAGGCCTCTATTCCCAGCTTCTT<br>GGCCTCCTCCTCGGTCGTCATGAACAGGCTAGTCCTCTGCCTTCCGCCCATGCTCCA<br>CATCATGTAGTAGACGACCAAGACAGT | none |
| SDSI 43 | ACAGTTCTCCTTCTTAGCTTCGTGAGAACGTTCAGCATAAAAGACGGTTTCACGGGC<br>CAAAGCCTAAGCGGCGTAACGGTGAAAGAAGGAGATACGGTTTTGGGCACGATTGA<br>CGACGGCGGGACGCTGGAGCTCACGAGGGGCACTCACACCTTGACTTTCGAGAAG<br>CCACATCATGTAGTAGACGACCAAGACAGT | none |
| SDSI 44 | ACAGTTCTCCTTCTTAGCTTCGTGAGAACCTGATGTTATAGAAGTCCGCAAGGACGG<br>CTCTGTCATCTCGCCCGAGGGTGGGAAATACTATCTCGGCGACATAAGCGGCCCGA<br>CACAAATTAGCATCAAGTTCAAGGCCGGCGCGGTGGGAACCCACGGCTTCACTATC<br>CACATCATGTAGTAGACGACCAAGACAGT | none |
| SDSI 45 | ACAGTTCTCCTTCTTAGCTTCGTGAGAACTCTCCCTCAACCTTCGCGGGGAGAACGG<br>CGCGGAGTACTGGACGGGCTACGCGGACGCGCTGGAAGACCTGTTGAAGAAAATCC<br>AGAGGCGGGAGGTGAGGGCATGAGAAGGTATTGTTACATCACGTGGGGATGGATCA<br>CACATCATGTAGTAGACGACCAAGACAGT | none |
| SDSI 46 | ACAGTTCTCCTTCTTAGCTTCGTGAGAACGAGCGCCGGGAGGTGAGGGCATGAGTG<br>AGGAATTGATGTTTGGTCGTGTCGTGGAGTATGTTCAGCATAGTTTCTACAAGAAACC<br>GTTTCCTCTTGGCAGTGAGCTCAAGAATGCAGTAGAGAAGGTTATGGAAACAGGACA<br>CATCATGTAGTAGACGACCAAGACAGT | none |
| SDSI 47 | ACAGTTCTCCTTCTTAGCTTCGTGAGAACAGGTCAGAGCCCACGTGGCAACTTTTGA<br>GGTTCTGACAAAAGACTATGTTCGTGAGAAATACAAAGACATCATAGAGTTCATGAGG<br>GAGAAAGGGACAGTATCGAGAAAGGAACTGCGGAAGAAGTTCTTCTTGCTTGCTCAC<br>ATCATGTAGTAGACGACCAAGACAGT | none |
| SDSI 48 | ACAGTTCTCCTTCTTAGCTTCGTGAGAACGTACCTCAAAATACAGAATCATATTTTACA<br>ATCGCTTGGAAATATTAATATCAACAATACGCAAGTCCAAATTAACGTCCCTGGCAAA<br>CAGGTGACAATTTATACCCACGAAATACTAGATAACGCCAAAAAGGCACTCGCACAT<br>CATGTAGTAGACGACCAAGACAGT | none |

*significant homology is defined as >75% sequence identity over >75% query cover

#### Sup Table 2. SDSI and Viral Read Percentages

| Mock CT | Viral reads % | Spike-in reads % |
|---|---|---|
| 20 | 99.56 | 0.18 |
| 25 | 99.19 | 0.38 |
| 30 | 98.47 | 1.11 |
| 35 | 99.65 | 3.17 |

#### Sup Table 3. ARTIC v3 Primers and Primers Spiked in at 2X

| Name | Pool | Sequence | Length | %GC | Spiked in at 2X |
|---|---|---|---|---|---|
| nCoV-2019_1_LEFT | nCoV-2019_1 | ACCAACCAACTTTCGATCTCTTGT | 24 | 41.7 | |
| nCoV-2019_1_RIGHT | nCoV-2019_1 | CATCTTTAAGATGTTGACGTGCCTC | 25 | 44.0 | |
| nCoV-2019_2_LEFT | nCoV-2019_2 | CTGTTTTACAGGTTCGCGACGT | 22 | 50.0 | |
| nCoV-2019_2_RIGHT | nCoV-2019_2 | TAAGGATCAGTGCCAAGCTCGT | 22 | 50.0 | |
| nCoV-2019_3_LEFT | nCoV-2019_1 | CGGTAATAAAGGAGCTGGTGGC | 22 | 54.6 | |
| nCoV-2019_3_RIGHT | nCoV-2019_1 | AAGGTGTCTGCAATTCATAGCTCT | 24 | 41.7 | |

| | | | | | |
|---|---|---|---|---|---|
| nCoV-2019_4_LEFT | nCoV-2019_2 | GGTGTATACTGCTGCCGTGAAC | 22 | 54.6 | |
| nCoV-2019_4_RIGHT | nCoV-2019_2 | CACAAGTAGTGGCACCTTCTTTAGT | 25 | 44.0 | |
| nCoV-2019_5_LEFT | nCoV-2019_1 | TGGTGAAACTTCATGGCAGACG | 22 | 50.0 | |
| nCoV-2019_5_RIGHT | nCoV-2019_1 | ATTGATGTTGACTTTCTCTTTTTGGAGT | 28 | 32.1 | |
| nCoV-2019_6_LEFT | nCoV-2019_2 | GGTGTTGTTGGAGAAGGTTCCG | 22 | 54.6 | |
| nCoV-2019_6_RIGHT | nCoV-2019_2 | TAGCGGCCTTCTGTAAAACACG | 22 | 50.0 | |
| nCoV-2019_7_LEFT_alt0 | nCoV-2019_1 | CATTTGCATCAGAGGCTGCTCG | 22 | 54.6 | X |
| nCoV-2019_7_RIGHT_alt5 | nCoV-2019_1 | AGGTGACAATTTGTCCACCGAC | 22 | 50.0 | X |
| nCoV-2019_8_LEFT | nCoV-2019_2 | AGAGTTTCTTAGAGACGGTTGGGA | 24 | 45.8 | |
| nCoV-2019_8_RIGHT | nCoV-2019_2 | GCTTCAACAGCTTCACTAGTAGGT | 24 | 45.8 | |
| nCoV-2019_9_LEFT_alt4 | nCoV-2019_1 | TTCCCACAGAAGTGTTAACAGAGG | 24 | 45.8 | X |
| nCoV-2019_9_RIGHT_alt2 | nCoV-2019_1 | GACAGCATCTGCCACAACACAG | 22 | 54.6 | X |
| nCoV-2019_10_LEFT | nCoV-2019_2 | TGAGAAGTGCTCTGCCTATACAGT | 24 | 45.8 | |
| nCoV-2019_10_RIGHT | nCoV-2019_2 | TCATCTAACCAATCTTCTTCTTGCTCT | 27 | 37.0 | |
| nCoV-2019_11_LEFT | nCoV-2019_1 | GGAATTTGGTGCCACTTCTGCT | 22 | 50.0 | |
| nCoV-2019_11_RIGHT | nCoV-2019_1 | TCATCAGATTCAACTTGCATGGCA | 24 | 41.7 | |
| nCoV-2019_12_LEFT | nCoV-2019_2 | AAACATGGAGGAGGTGTTGCAG | 22 | 50.0 | X |
| nCoV-2019_12_RIGHT | nCoV-2019_2 | TTCACTCTTCATTTCCAAAAAGCTTGA | 27 | 33.3 | X |
| nCoV-2019_13_LEFT | nCoV-2019_1 | TCGCACAAATGTCTACTTAGCTGT | 24 | 41.7 | |
| nCoV-2019_13_RIGHT | nCoV-2019_1 | ACCACAGCAGTTAAAACACCCT | 22 | 45.5 | |
| nCoV-2019_14_LEFT_alt4 | nCoV-2019_2 | TGGCAATCTTCATCCAGATTCTGC | 24 | 45.8 | X |
| nCoV-2019_14_RIGHT_alt2 | nCoV-2019_2 | TGCGTGTTTCTTCTGCATGTGC | 22 | 50.0 | X |
| nCoV-2019_15_LEFT_alt1 | nCoV-2019_1 | AGTGCTTAAAAAGTGTAAAAGTGCCT | 26 | 34.6 | X |
| nCoV-2019_15_RIGHT_alt3 | nCoV-2019_1 | ACTGTAGCTGGCACTTTGAGAGA | 23 | 47.8 | X |
| nCoV-2019_16_LEFT | nCoV-2019_2 | AATTTGGAAGAAGCTGCTCGGT | 22 | 45.5 | |
| nCoV-2019_16_RIGHT | nCoV-2019_2 | CACAACTTGCGTGTGGAGGTTA | 22 | 50.0 | |
| nCoV-2019_17_LEFT | nCoV-2019_1 | CTTCTTTCTTTGAGAGAAGTGAGGACT | 27 | 40.7 | X |
| nCoV-2019_17_RIGHT | nCoV-2019_1 | TTTGTTGGAGTGTTAACAATGCAGT | 25 | 36.0 | X |
| nCoV-2019_18_LEFT_alt2 | nCoV-2019_2 | ACTTCTATTAAATGGGCAGATAACAACTGT | 30 | 33.3 | X |
| nCoV-2019_18_RIGHT_alt1 | nCoV-2019_2 | GCTTGTTTACCACACGTACAAGG | 23 | 47.8 | X |
| nCoV-2019_19_LEFT | nCoV-2019_1 | GCTGTTATGTACATGGGCACACT | 23 | 47.8 | |
| nCoV-2019_19_RIGHT | nCoV-2019_1 | TGTCCAACTTAGGGTCAATTTCTGT | 25 | 40.0 | |
| nCoV-2019_20_LEFT | nCoV-2019_2 | ACAAAGAAAACAGTTACACAACAACCA | 27 | 33.3 | |
| nCoV-2019_20_RIGHT | nCoV-2019_2 | ACGTGGCTTTATTAGTTGCATTGTT | 25 | 36.0 | |
| nCoV-2019_21_LEFT_alt2 | nCoV-2019_1 | GGCTATTGATTATAAACACTACACACCCT | 29 | 37.9 | X |
| nCoV-2019_21_RIGHT_alt0 | nCoV-2019_1 | GATCTGTGTGGCCAACCTCTTC | 22 | 54.6 | X |
| nCoV-2019_22_LEFT | nCoV-2019_2 | ACTACCGAAGTTGTAGGAGACATTATACT | 29 | 37.9 | |

| | | | | | |
|---|---|---|---|---|---|
| nCoV-2019_22_RIGHT | nCoV-2019_2 | ACAGTATTCTTTGCTATAGTAGTCGGC | 27 | 40.7 | |
| nCoV-2019_23_LEFT | nCoV-2019_1 | ACAACTACTAACATAGTTACACGGTGT | 27 | 37.0 | |
| nCoV-2019_23_RIGHT | nCoV-2019_1 | ACCAGTACAGTAGGTTGCAATAGTG | 25 | 44.0 | |
| nCoV-2019_24_LEFT | nCoV-2019_2 | AGGCATGCCTTCTTACTGTACTG | 23 | 47.8 | X |
| nCoV-2019_24_RIGHT | nCoV-2019_2 | ACATTCTAACCATAGCTGAAATCGGG | 26 | 42.3 | X |
| nCoV-2019_25_LEFT | nCoV-2019_1 | GCAATTGTTTTTCAGCTATTTTGCAGT | 27 | 33.3 | |
| nCoV-2019_25_RIGHT | nCoV-2019_1 | ACTGTAGTGACAAGTCTCTCGCA | 23 | 47.8 | |
| nCoV-2019_26_LEFT | nCoV-2019_2 | TTGTGATACATTCTGTGCTGGTAGT | 25 | 40.0 | |
| nCoV-2019_26_RIGHT | nCoV-2019_2 | TCCGCACTATCACCAACATCAG | 22 | 50.0 | |
| nCoV-2019_27_LEFT | nCoV-2019_1 | ACTACAGTCAGCTTATGTGTCAACC | 25 | 44.0 | |
| nCoV-2019_27_RIGHT | nCoV-2019_1 | AATACAAGCACCAAGGTCACGG | 22 | 50.0 | |
| nCoV-2019_28_LEFT | nCoV-2019_2 | ACATAGAAGTTACTGGCGATAGTTGT | 26 | 38.5 | |
| nCoV-2019_28_RIGHT | nCoV-2019_2 | TGTTTAGACATGACATGAACAGGTGT | 26 | 38.5 | |
| nCoV-2019_29_LEFT | nCoV-2019_1 | ACTTGTGTTCCTTTTTGTTGCTGC | 24 | 41.7 | |
| nCoV-2019_29_RIGHT | nCoV-2019_1 | AGTGTACTCTATAAGTTTTGATGGTGTGT | 29 | 34.5 | |
| nCoV-2019_30_LEFT | nCoV-2019_2 | GCACAACTAATGGTGACTTTTTGCA | 25 | 40.0 | |
| nCoV-2019_30_RIGHT | nCoV-2019_2 | ACCACTAGTAGATACACAAACACCAG | 26 | 42.3 | |
| nCoV-2019_31_LEFT | nCoV-2019_1 | TTCTGAGTACTGTAGGCACGGC | 22 | 54.6 | |
| nCoV-2019_31_RIGHT | nCoV-2019_1 | ACAGAATAAACACCAGGTAAGAATGAGT | 28 | 35.7 | |
| nCoV-2019_32_LEFT | nCoV-2019_2 | TGGTGAATACAGTCATGTAGTTGCC | 25 | 44.0 | |
| nCoV-2019_32_RIGHT | nCoV-2019_2 | AGCACATCACTACGCAACTTTAGA | 24 | 41.7 | |
| nCoV-2019_33_LEFT | nCoV-2019_1 | ACTTTTGAAGAAGCTGCGCTGT | 22 | 45.5 | X |
| nCoV-2019_33_RIGHT | nCoV-2019_1 | TGGACAGTAAACTACGTCATCAAGC | 25 | 44.0 | X |
| nCoV-2019_34_LEFT | nCoV-2019_2 | TCCCATCTGGTAAAGTTGAGGGT | 23 | 47.8 | |
| nCoV-2019_34_RIGHT | nCoV-2019_2 | AGTGAAATTGGGCCTCATAGCA | 22 | 45.5 | |
| nCoV-2019_35_LEFT | nCoV-2019_1 | TGTTCGCATTCAACCAGGACAG | 22 | 50.0 | |
| nCoV-2019_35_RIGHT | nCoV-2019_1 | ACTTCATAGCCACAAGGTTAAAGTCA | 26 | 38.5 | |
| nCoV-2019_36_LEFT | nCoV-2019_2 | TTAGCTTGGTTGTACGCTGCTG | 22 | 50.0 | |
| nCoV-2019_36_RIGHT | nCoV-2019_2 | GAACAAAGACCATTGAGTACTCTGGA | 26 | 42.3 | |
| nCoV-2019_37_LEFT | nCoV-2019_1 | ACACACCACTGGTTGTTACTCAC | 23 | 47.8 | |
| nCoV-2019_37_RIGHT | nCoV-2019_1 | GTCCACACTCTCCTAGCACCAT | 22 | 54.6 | |
| nCoV-2019_38_LEFT | nCoV-2019_2 | ACTGTGTTATGTATGCATCAGCTGT | 25 | 40.0 | |
| nCoV-2019_38_RIGHT | nCoV-2019_2 | CACCAAGAGTCAGTCTAAAGTAGCG | 25 | 48.0 | |
| nCoV-2019_39_LEFT | nCoV-2019_1 | AGTATTGCCCTATTTTCTTCATAACTGGT | 29 | 34.5 | |
| nCoV-2019_39_RIGHT | nCoV-2019_1 | TGTAACTGGACACATTGAGCCC | 22 | 50.0 | |
| nCoV-2019_40_LEFT | nCoV-2019_2 | TGCACATCAGTAGTCTTACTCTCAGT | 26 | 42.3 | |
| nCoV-2019_40_RIGHT | nCoV-2019_2 | CATGGCTGCATCACGGTCAAAT | 22 | 50.0 | |

| | | | | | |
|---|---|---|---|---|---|
| nCoV-2019_41_LEFT | nCoV-2019_1 | GTTCCCTTCCATCATATGCAGCT | 23 | 47.8 | |
| nCoV-2019_41_RIGHT | nCoV-2019_1 | TGGTATGACAACCATTAGTTTGGCT | 25 | 40.0 | |
| nCoV-2019_42_LEFT | nCoV-2019_2 | TGCAAGAGATGGTTGTGTTCCC | 22 | 50.0 | |
| nCoV-2019_42_RIGHT | nCoV-2019_2 | CCTACCTCCCTTTGTTGTGTTGT | 23 | 47.8 | |
| nCoV-2019_43_LEFT | nCoV-2019_1 | TACGACAGATGTCTTGTGCTGC | 22 | 50.0 | |
| nCoV-2019_43_RIGHT | nCoV-2019_1 | AGCAGCATCTACAGCAAAAGCA | 22 | 45.5 | |
| nCoV-2019_44_LEFT_alt3 | nCoV-2019_2 | CCACAGTACGTCTACAAGCTGG | 22 | 54.6 | |
| nCoV-2019_44_RIGHT_alt0 | nCoV-2019_2 | CGCAGACGGTACAGACTGTGTT | 22 | 54.6 | |
| nCoV-2019_45_LEFT_alt2 | nCoV-2019_1 | AGTATGTACAAATACCTACAACTTGTGCT | 29 | 34.5 | X |
| nCoV-2019_45_RIGHT_alt7 | nCoV-2019_1 | TTCATGTTGGTAGTTAGAGAAAGTGTGTC | 29 | 37.9 | X |
| nCoV-2019_46_LEFT_alt1 | nCoV-2019_2 | CGCTTCCAAGAAAAGGACGAAGA | 23 | 47.8 | |
| nCoV-2019_46_RIGHT_alt2 | nCoV-2019_2 | CACGTTCACCTAAGTTGGCGTAT | 23 | 47.8 | |
| nCoV-2019_47_LEFT | nCoV-2019_1 | AGGACTGGTATGATTTTGTAGAAAACCC | 28 | 39.3 | |
| nCoV-2019_47_RIGHT | nCoV-2019_1 | AATAACGGTCAAAGAGTTTTAACCTCTC | 28 | 35.7 | |
| nCoV-2019_48_LEFT | nCoV-2019_2 | TGTTGACACTGACTTAACAAAGCCT | 25 | 40.0 | |
| nCoV-2019_48_RIGHT | nCoV-2019_2 | TAGATTACCAGAAGCAGCGTGC | 22 | 50.0 | |
| nCoV-2019_49_LEFT | nCoV-2019_1 | AGGAATTACTTGTGTATGCTGCTGA | 25 | 40.0 | |
| nCoV-2019_49_RIGHT | nCoV-2019_1 | TGACGATGACTTGGTTAGCATTAATACA | 28 | 35.7 | |
| nCoV-2019_50_LEFT | nCoV-2019_2 | GTTGATAAGTACTTTGATTGTTACGATGGT | 30 | 33.3 | |
| nCoV-2019_50_RIGHT | nCoV-2019_2 | TAACATGTTGTGCCAACCACCA | 22 | 45.5 | |
| nCoV-2019_51_LEFT | nCoV-2019_1 | TCAATAGCCGCCACTAGAGGAG | 22 | 54.6 | |
| nCoV-2019_51_RIGHT | nCoV-2019_1 | AGTGCATTAACATTGGCCGTGA | 22 | 45.5 | |
| nCoV-2019_52_LEFT | nCoV-2019_2 | CATCAGGAGATGCCACAACTGC | 22 | 54.6 | |
| nCoV-2019_52_RIGHT | nCoV-2019_2 | GTTGAGAGCAAAATTCATGAGGTCC | 25 | 44.0 | |
| nCoV-2019_53_LEFT | nCoV-2019_1 | AGCAAAATGTTGGACTGAGACTGA | 24 | 41.7 | |
| nCoV-2019_53_RIGHT | nCoV-2019_1 | AGCCTCATAAAACTCAGGTTCCC | 23 | 47.8 | |
| nCoV-2019_54_LEFT | nCoV-2019_2 | TGAGTTAACAGGACACATGTTAGACA | 26 | 38.5 | |
| nCoV-2019_54_RIGHT | nCoV-2019_2 | AACCAAAAACTTGTCCATTAGCACA | 25 | 36.0 | |
| nCoV-2019_55_LEFT | nCoV-2019_1 | ACTCAACTTTACTTAGGAGGTATGAGCT | 28 | 39.3 | |
| nCoV-2019_55_RIGHT | nCoV-2019_1 | GGTGTACTCTCCTATTTGTACTTTACTGT | 29 | 37.9 | |
| nCoV-2019_56_LEFT | nCoV-2019_2 | ACCTAGACCACCACTTAACCGA | 22 | 50.0 | |
| nCoV-2019_56_RIGHT | nCoV-2019_2 | ACACTATGCGAGCAGAAGGGTA | 22 | 50.0 | |
| nCoV-2019_57_LEFT | nCoV-2019_1 | ATTCTACACTCCAGGGACCACC | 22 | 54.6 | |
| nCoV-2019_57_RIGHT | nCoV-2019_1 | GTAATTGAGCAGGGTCGCCAAT | 22 | 50.0 | |
| nCoV-2019_58_LEFT | nCoV-2019_2 | TGATTTGAGTGTTGTCAATGCCAGA | 25 | 40.0 | |
| nCoV-2019_58_RIGHT | nCoV-2019_2 | CTTTTCTCCAAGCAGGGTTACGT | 23 | 47.8 | |
| nCoV-2019_59_LEFT | nCoV-2019_1 | TCACGCATGATGTTTCATCTGCA | 23 | 43.5 | |

| | | | | | |
|---|---|---|---|---|---|
| nCoV-2019_59_RIGHT | nCoV-2019_1 | AAGAGTCCTGTTACATTTTCAGCTTG | 26 | 38.5 | |
| nCoV-2019_60_LEFT | nCoV-2019_2 | TGATAGAGACCTTTATGACAAGTTGCA | 27 | 37.0 | |
| nCoV-2019_60_RIGHT | nCoV-2019_2 | GGTACCAACAGCTTCTCTAGTAGC | 24 | 50.0 | |
| nCoV-2019_61_LEFT | nCoV-2019_1 | TGTTTATCACCCGCGAAGAAGC | 22 | 50.0 | |
| nCoV-2019_61_RIGHT | nCoV-2019_1 | ATCACATAGACAACAGGTGCGC | 22 | 50.0 | |
| nCoV-2019_62_LEFT | nCoV-2019_2 | GGCACATGGCTTTGAGTTGACA | 22 | 50.0 | |
| nCoV-2019_62_RIGHT | nCoV-2019_2 | GTTGAACCTTTCTACAAGCCGC | 22 | 50.0 | |
| nCoV-2019_63_LEFT | nCoV-2019_1 | TGTTAAGCGTGTTGACTGGACT | 22 | 45.5 | |
| nCoV-2019_63_RIGHT | nCoV-2019_1 | ACAAACTGCCACCATCACAACC | 22 | 50.0 | |
| nCoV-2019_64_LEFT | nCoV-2019_2 | TCGATAGATATCCTGCTAATTCCATTGT | 28 | 35.7 | X |
| nCoV-2019_64_RIGHT | nCoV-2019_2 | AGTCTTGTAAAAGTGTTCCAGAGGT | 25 | 40.0 | X |
| nCoV-2019_65_LEFT | nCoV-2019_1 | GCTGGCTTTAGCTTGTGGGTTT | 22 | 50.0 | |
| nCoV-2019_65_RIGHT | nCoV-2019_1 | TGTCAGTCATAGAACAAACACCAATAGT | 28 | 35.7 | |
| nCoV-2019_66_LEFT | nCoV-2019_2 | GGGTGTGGACATTGCTGCTAAT | 22 | 50.0 | X |
| nCoV-2019_66_RIGHT | nCoV-2019_2 | TCAATTTCCATTTGACTCCTGGGT | 24 | 41.7 | X |
| nCoV-2019_67_LEFT | nCoV-2019_1 | GTTGTCCAACAATTACCTGAAACTTACT | 28 | 35.7 | X |
| nCoV-2019_67_RIGHT | nCoV-2019_1 | CAACCTTAGAAACTACAGATAAATCTTGGG | 30 | 36.7 | X |
| nCoV-2019_68_LEFT | nCoV-2019_2 | ACAGGTTCATCTAAGTGTGTGTGT | 24 | 41.7 | |
| nCoV-2019_68_RIGHT | nCoV-2019_2 | CTCCTTTATCAGAACCAGCACCA | 23 | 47.8 | |
| nCoV-2019_69_LEFT | nCoV-2019_1 | TGTCGCAAAATATACTCAACTGTGTCA | 27 | 37.0 | |
| nCoV-2019_69_RIGHT | nCoV-2019_1 | TCTTTATAGCCACGGAACCTCCA | 23 | 47.8 | |
| nCoV-2019_70_LEFT | nCoV-2019_2 | ACAAAAGAAAATGACTCTAAAGAGGGTTT | 29 | 31.0 | X |
| nCoV-2019_70_RIGHT | nCoV-2019_2 | TGACCTTCTTTTAAAGACATAACAGCAG | 28 | 35.7 | X |
| nCoV-2019_71_LEFT | nCoV-2019_1 | ACAAATCCAATTCAGTTGTCTTCCTATTC | 29 | 34.5 | X |
| nCoV-2019_71_RIGHT | nCoV-2019_1 | TGGAAAAGAAAGGTAAGAACAAGTCCT | 27 | 37.0 | X |
| nCoV-2019_72_LEFT | nCoV-2019_2 | ACACGTGGTGTTTATTACCCTGAC | 24 | 45.8 | |
| nCoV-2019_72_RIGHT | nCoV-2019_2 | ACTCTGAACTCACTTTCCATCCAAC | 25 | 44.0 | |
| nCoV-2019_73_LEFT | nCoV-2019_1 | CAATTTTGTAATGATCCATTTTTGGGTGT | 29 | 31.0 | |
| nCoV-2019_73_RIGHT | nCoV-2019_1 | CACCAGCTGTCCAACCTGAAGA | 22 | 54.6 | |
| nCoV-2019_74_LEFT | nCoV-2019_2 | ACATCACTAGGTTTCAAACTTTACTTGC | 28 | 35.7 | |
| nCoV-2019_74_RIGHT | nCoV-2019_2 | GCAACACAGTTGCTGATTCTCTTC | 24 | 45.8 | |
| nCoV-2019_75_LEFT | nCoV-2019_1 | AGAGTCCAACCAACAGAATCTATTGT | 26 | 38.5 | |
| nCoV-2019_75_RIGHT | nCoV-2019_1 | ACCACCAACCTTAGAATCAAGATTGT | 26 | 38.5 | |
| nCoV-2019_76_LEFT_alt3 | nCoV-2019_2 | GGGCAAACTGGAAAGATTGCTGA | 23 | 47.8 | X |
| nCoV-2019_76_RIGHT_alt0 | nCoV-2019_2 | ACCTGTGCCTGTTAAACCATTGA | 23 | 43.5 | X |
| nCoV-2019_77_LEFT | nCoV-2019_1 | CCAGCAACTGTTTGTGGACCTA | 22 | 50.0 | |
| nCoV-2019_77_RIGHT | nCoV-2019_1 | CAGCCCCTATTAAACAGCCTGC | 22 | 54.6 | |

| | | | | | |
|---|---|---|---|---|---|
| nCoV-2019_78_LEFT | nCoV-2019_2 | CAACTTACTCCTACTTGGCGTGT | 23 | 47.8 | |
| nCoV-2019_78_RIGHT | nCoV-2019_2 | TGTGTACAAAAACTGCCATATTGCA | 25 | 36.0 | |
| nCoV-2019_79_LEFT | nCoV-2019_1 | GTGGTGATTCAACTGAATGCAGC | 23 | 47.8 | X |
| nCoV-2019_79_RIGHT | nCoV-2019_1 | CATTTCATCTGTGAGCAAAGGTGG | 24 | 45.8 | X |
| nCoV-2019_80_LEFT | nCoV-2019_2 | TTGCCTTGGTGATATTGCTGCT | 22 | 45.5 | X |
| nCoV-2019_80_RIGHT | nCoV-2019_2 | TGGAGCTAAGTTGTTTAACAAGCG | 24 | 41.7 | X |
| nCoV-2019_81_LEFT | nCoV-2019_1 | GCACTTGGAAAACTTCAAGATGTGG | 25 | 44.0 | |
| nCoV-2019_81_RIGHT | nCoV-2019_1 | GTGAAGTTCTTTTCTTGTGCAGGG | 24 | 45.8 | |
| nCoV-2019_82_LEFT | nCoV-2019_2 | GGGCTATCATCTTATGTCCTTCCCT | 25 | 48.0 | |
| nCoV-2019_82_RIGHT | nCoV-2019_2 | TGCCAGAGATGTCACCTAAATCAA | 24 | 41.7 | |
| nCoV-2019_83_LEFT | nCoV-2019_1 | TCCTTTGCAACCTGAATTAGACTCA | 25 | 40.0 | |
| nCoV-2019_83_RIGHT | nCoV-2019_1 | TTTGACTCCTTTGAGCACTGGC | 22 | 50.0 | |
| nCoV-2019_84_LEFT | nCoV-2019_2 | TGCTGTAGTTGTCTCAAGGGCT | 22 | 50.0 | |
| nCoV-2019_84_RIGHT | nCoV-2019_2 | AGGTGTGAGTAAACTGTTACAAACAAC | 27 | 37.0 | |
| nCoV-2019_85_LEFT | nCoV-2019_1 | ACTAGCACTCTCCAAGGGTGTT | 22 | 50.0 | |
| nCoV-2019_85_RIGHT | nCoV-2019_1 | ACACAGTCTTTTACTCCAGATTCCC | 25 | 44.0 | |
| nCoV-2019_86_LEFT | nCoV-2019_2 | TCAGGTGATGGCACAACAAGTC | 22 | 50.0 | |
| nCoV-2019_86_RIGHT | nCoV-2019_2 | ACGAAAGCAAGAAAAAGAAGTACGC | 25 | 40.0 | |
| nCoV-2019_87_LEFT | nCoV-2019_1 | CGACTACTAGCGTGCCTTTGTA | 22 | 50.0 | |
| nCoV-2019_87_RIGHT | nCoV-2019_1 | ACTAGGTTCCATTGTTCAAGGAGC | 24 | 45.8 | |
| nCoV-2019_88_LEFT | nCoV-2019_2 | CCATGGCAGATTCCAACGGTAC | 22 | 54.6 | |
| nCoV-2019_88_RIGHT | nCoV-2019_2 | TGGTCAGAATAGTGCCATGGAGT | 23 | 47.8 | |
| nCoV-2019_89_LEFT_alt2 | nCoV-2019_1 | CGCGTTCCATGTGGTCATTCAA | 22 | 50.0 | |
| nCoV-2019_89_RIGHT_alt4 | nCoV-2019_1 | ACGAGATGAAACATCTGTTGTCACT | 25 | 40.0 | |
| nCoV-2019_90_LEFT | nCoV-2019_2 | ACACAGACCATTCCAGTAGCAGT | 23 | 47.8 | |
| nCoV-2019_90_RIGHT | nCoV-2019_2 | TGAAATGGTGAATTGCCCTCGT | 22 | 45.5 | |
| nCoV-2019_91_LEFT | nCoV-2019_1 | TCACTACCAAGAGTGTGTTAGAGGT | 25 | 44.0 | X |
| nCoV-2019_91_RIGHT | nCoV-2019_1 | TTCAAGTGAGAACCAAAAGATAATAAGCA | 29 | 31.0 | X |
| nCoV-2019_92_LEFT | nCoV-2019_2 | TTTGTGCTTTTTAGCCTTTCTGCT | 24 | 37.5 | |
| nCoV-2019_92_RIGHT | nCoV-2019_2 | AGGTTCCTGGCAATTAATTGTAAAAGG | 27 | 37.0 | |
| nCoV-2019_93_LEFT | nCoV-2019_1 | TGAGGCTGGTTCTAAATCACCCA | 23 | 47.8 | |
| nCoV-2019_93_RIGHT | nCoV-2019_1 | AGGTCTTCCTTGCCATGTTGAG | 22 | 50.0 | |
| nCoV-2019_94_LEFT | nCoV-2019_2 | GGCCCCAAGGTTTACCCAATAA | 22 | 50.0 | |
| nCoV-2019_94_RIGHT | nCoV-2019_2 | TTTGGCAATGTTGTTCCTTGAGG | 23 | 43.5 | |
| nCoV-2019_95_LEFT | nCoV-2019_1 | TGAGGGAGCCTTGAATACACCA | 22 | 50.0 | |
| nCoV-2019_95_RIGHT | nCoV-2019_1 | CAGTACGTTTTTGCCGAGGCTT | 22 | 50.0 | |
| nCoV-2019_96_LEFT | nCoV-2019_2 | GCCAACAACAACAAGGCCAAAC | 22 | 50.0 | |

| | | | | |
|---|---|---|---|---|
| nCoV-2019_96_RIGHT | nCoV-2019_2 | TAGGCTCTGTTGGTGGGAATGT | 22 | 50.0 |
| nCoV-2019_97_LEFT | nCoV-2019_1 | TGGATGACAAAGATCCAAATTTCAAAGA | 28 | 32.1 |
| nCoV-2019_97_RIGHT | nCoV-2019_1 | ACACACTGATTAAAGATTGCTATGTGAG | 28 | 35.7 |
| nCoV-2019_98_LEFT | nCoV-2019_2 | AACAATTGCAACAATCCATGAGCA | 24 | 37.5 |
| nCoV-2019_98_RIGHT | nCoV-2019_2 | TTCTCCTAAGAAGCTATTAAAATCACATGG | 30 | 33.3 |

### Sup Table 4. Time and Cost Comparison of FLEX vs XT

| Library Prep Kit | Cost Per Sample ($) | Time (hrs) |
|---|---|---|
| Illumina DNA Flex | 45.96 | 10 |
| Illumina Nextera XT | 64.43 | 13.5 |

### Sup Table 5. Cost of SDSI+ARTIC

| Processing Step | Reagent | Vendor | Item Number | Cost (dollars) | Number of Reactions | Cost per Reaction |
|---|---|---|---|---|---|---|
| Biosample Extraction | MagMAX™ mirVana™ Total RNA Isolation Kit | Thermo Fisher Scientific | A27828 | 495 | 96 | 5.16 |
| | SSIV RT master mix | Thermo Fisher Scientific | 18090050 | 383 | 50 | 7.66 |
| cDNA Synthesis | Random hexamers (50ng/ul) | Thermo Fisher Scientific | N808127 | 91 | 100 | 0.91 |
| | dNTPs (10nM) | Thermo Fisher Scientific | 18427-013 | 99 | 100 | 0.99 |
| | 5x RT buffer | Thermo Fisher Scientific | 18090050 | x | x | x |
| | DTT (100mM) | Thermo Fisher Scientific | 18090050 | x | x | x |
| | Superase rnase inhibitor | Thermo Fisher Scientific | 10777-019 | 188 | 125 | 1.50 |
| ARTIC PCR | Q5 Hot Start High-Fidelity 2X Master Mix | New England BioLabs | M0494L | 845 | 500 | 1.69 |
| | Artic Primers Pool#1 and Pool#2 | IDT | | 30 | 500 | 0.06 |
| Spike-ins | Spike in Primers (Forward/Reverse) | IDT | | 500 | 1000000 | 0.00 |
| | Spike-in targets n=96 | IDT | | 5821 | 1000000 | 0.01 |
| Post Artic Pooling Quantification | Qubit™ dsDNA HS Assay Kit | Thermo Fisher Scientific | Q32854 | 308 | 500 | 0.62 |
| Library Construction | Nextera DNA flex Library Prep (n=96) | Illumina | 20018705 | 4153 | 190 | 21.86 |
| | Nextera index UD Set A (n=96) | Illumina | 20027213 | 672 | 384 | 1.75 |
| Library Quantification | High Sensitivity D1000 ScreenTape | Agilent | 5067-5584 | 362 | 112 | 3.23 |
| | High Sensitivity D1000 Sample Buffer | Agilent | 5067-5603 | 59.14 | 112 | 0.53 |
| **TOTAL:** | | | | | | 45.96 |
